# Molecular Basis for Impacts of DSIF on the Dynamics of RNA Polymerase II Elongation Complex

**DOI:** 10.1101/2025.08.09.669504

**Authors:** Weththasinghage D. Amith, Brandon M. Bogart, Bercem Dutagaci

**Affiliations:** Department of Molecular and Cell Biology, University of California, Merced, 5200 North Lake Rd. Merced, CA 95343, USA; Health Sciences Research Institute, University of California, Merced, 5200 North Lake Rd. Merced, CA 95343, USA

## Abstract

Transcription elongation is a highly regulated process involving elongation factors associated with RNA polymerase II (Pol II). DRB sensitivity-inducing factor (DSIF) is an elongation factor known to have multiple roles in transcription elongation. Although studies resolved the structures of elongation complexes with DSIF, little is known about the impacts of DSIF on the dynamics of the elongation complex at the molecular level. Here, we used molecular dynamics simulations to elucidate the effects of DSIF on the dynamics and structure of Pol II and upstream nucleic acids, thereby gaining a mechanistic understanding of its role in transcription elongation. We determined three major sites of impact by DSIF, including the upstream nucleic acids, Pol II clamp, and active site, which potentially contribute to its role in transcription processivity. Our results showed that DSIF affects the dynamics of upstream DNA and RNA at the exit sites, preventing the unwinding of DNA and the folding of RNA. In addition, our results suggest that DSIF regulates the motion of the Pol II clamp to potentially maintain a proper size at the central cleft. We also observed a more dynamic active site and increased interactions between active site domains. Based on correlated motion analysis, we proposed that the impacts of DSIF on the active site have an allosteric nature that takes place through the collective motions of the Pol II clamp and nucleic acids.

## 1. INTRODUCTION

RNA polymerase II (Pol II) is the central enzyme in transcription of all mRNAs and many noncoding RNAs in eukaryotes [1]. Elongation factors accommodate Pol II in transition from the initiation stage of transcription to the stages of promoter-proximal pausing and subsequently elongation [2–4]. Elongation rate and processivity are highly regulated by the collective functions of various elongation factors, including DRB sensitivity-inducing factor (DSIF, also known as Spt4/5), Spt6, polymerase-associated factor 1 complex (PAF), and positive transcription elongation factor (P-TEFb) [5–17]. DSIF is a highly conserved elongation factor in all three domains of life and is formed by the association of Spt4 and Spt5 factors in Eukarya and Archaea and NusG (Spt5 homolog) in Bacteria. DSIF has important roles in promoter-proximal pausing [8,18–21], transcription processivity [6,10,16,22–24], and nucleosomal access [7,25–28], while the molecular mechanisms of the roles of DSIF in transcription elongation remain to be resolved.

Transcription processivity describes the process of transcription elongation in which Pol II remains activated and able to maintain elongation without interruptions [29]. Elongation factors are proposed to maintain transcription processivity by preventing the formation of paused/arrested states of Pol II or releasing Pol II from those states [19,30]. DSIF is one of the elongation factors that was shown to enhance processivity. Earlier studies demonstrated that DSIF stimulates Pol II elongation activity [16,31,32] and reduces or prevents the formation of arrested/paused states [6,33]. There are various hypotheses on the mechanisms of the DSIF function in maintaining transcription processivity. Some studies suggested that DSIF stabilized a closed clamp conformation, which is necessary for maintaining active elongation [24,34]. More recently, in support of this hypothesis, another study demonstrated the formation of a closer clamp in the presence of Spt4/5 in an archaeal RNA polymerase using the cryo-electron microscopy (cryo-EM) technique [35]. Other studies focused on the interactions between Spt4/5 with DNA and suggested that Spt4/5 increases processivity by encompassing a DNA clamp, which prevents DNA from disengaging or forming secondary structures [6,36]. More recently, structural studies using Cryo-EM techniques showed that DSIF forms DNA and RNA clamps at the exit sites, supporting the importance of DSIF interactions with upstream nucleic acids in transcription processivity [12,22,23,28]. Furthermore, studies suggest an allosteric effect of DSIF on active site dynamics [6,24,31,33,37]. The allosteric function of DSIF was proposed to take place in two potential mechanisms, one is through the impacts of DSIF on the structure and dynamics of DNA and RNA [6,33,37], and the other is through the interactions of DSIF with the Pol II clamp domain [6,24,31]. In both mechanisms, DSIF is potentially impacting the structure and dynamics of the trigger loop (TL) and bridge helix (BH), both located near the active site and may help maintain processivity by preventing pausing and arrest [6] and promoting forward translocation [33,37].

Previous computational studies on transcription elongation predominantly focused on the Pol II active site in the absence of elongation factors [38–47]. These studies used molecular dynamics (MD) simulations, machine learning and Markov state modeling (MSM) and provided important insights into the roles of TL and BH in positioning the incoming NTP, transcription fidelity, translocation and backtracking. After cryo-EM structures of Pol II with elongation factors have been resolved by multiple research groups [12,22,23,25,28,48,49], our lab has started to study the Pol II-DSIF complex [50,51]. We previously showed that the interactions between DSIF and upstream nucleic acids are strongly sustained throughout MD simulations in the complete Pol II-DSIF complex and isolated Spt5-upstream nucleic acid systems [50], supporting the importance of DSIF’s impact on nucleic acid dynamics for its role in transcription elongation. Motivated by our earlier work, here we studied the Pol II elongation complex in the presence and absence of DSIF to investigate the impacts of DSIF on the structure and dynamics of the Pol II elongation complex. Our simulations revealed three important sites of impact of DSIF on the Pol II complex, which are upstream nucleic acids, active site domains, and the Rpb1-clamp domain. DSIF has a strong impact on upstream nucleic acid dynamics to prevent unwinding and secondary structure formation. For the active site domains, we observed increased dynamics, especially for TL. In addition, the interactions between TL and BH increased in the presence of DSIF, and we propose that these interactions may play an important role in the TL opening/closing motion. The impacts of DSIF on the clamp motion were more subtle, probably due to the difficulty in capturing globular motions with the simulation time scale. We observed that the distances of the clamp to the jaw and protrusion domains across the central cleft slightly increased. We propose that this increase reflects DSIF’s role in ensuring proper clamp motion rather than closing the clamp further. Overall, based on our results, we suggest that these three impact points may work together to maintain the processivity of transcription.

## 2. METHODS

### 2.1. System Preparation

We prepared three systems of the Pol II complex for this study. Two of the systems were prepared using the published structure with PDB ID: 5OIK [22]. In this structure, Pol II is from *Bos taurus* and DSIF is from *Homo sapiens*. For the first system, we used the complete complex with DSIF, which is referred to as +DSIF. For the second system, we removed DSIF (Spt4/5) from the complex, which is referred to as -DSIF. In addition, we prepared another system without DSIF using the published Pol II complex structure (PDB ID: 5FLM) [52] and referred it as -DSIF_5flm. In this structure, Pol II is also from *Bos taurus,* and the two structures are similar, without any significant conformational differences (Table S1). We mostly discussed the results for +DSIF and -DSIF systems in the paper, however, we also presented the results of the -DSIF_5flm system and compared it with the -DSIF system. We modeled the missing residues of the TL using the MODELLER program [53]. The histidine at 1108 of Rpb1 was protonated, since it is the equivalent of H1085 in Pol II from *Saccharomyces cerevisiae*, which was suggested to be protonated by an earlier study [40]. Since the initial structure with PDB ID: 5FLM has shorter DNA and RNA at the exit sites, for the -DSIF_5flm system, we modeled upstream DNA and nascent RNA based on the structure with PDB ID: 5OIK by superimposing the two structures nucleic acid constructs. The systems were solvated in a cubic box by using at least a 9 Å cutoff from the edges of the simulation boxes. This resulted in simulation systems with system sizes of 180.2, 173.2, and 174.8 Å and numbers of atoms of 560,945, 497,751, and 514,233 for the +DSIF, -DSIF and -DSIF_5flm, respectively. We neutralize the systems by adding 116, 110 and 114 Na^+^ ions in +DSIF, -DSIF and -DSIF_5flm systems, respectively. System preparation was performed by using the MMTSB package [54] together with the CHARMM software version 45b2 [55].

### 2.2. MD simulations

The systems were minimized using 5,000 steps of energy minimization. Systems were then equilibrated for 1.6 ns while increasing the temperature from 100 to 300K and using constraints with a force constant of 400 and 40 kJ/mol/nm^2^ for the heavy atoms of the backbone and sidechain of proteins, respectively. After equilibration, we performed 500 ns of MD simulations for five replicates for +DSIF and -DSIF systems, which makes a total of 5 μs simulations. For the -DSIF_5flm system, we performed 250 ns of MD simulations for five replicates. Periodic boundary conditions and Particle Mesh Ewald algorithm [56,57] were applied for the calculation of long-range electrostatic interactions. Lennard-Jones interactions were switched between 10 and 12 Å. Bonds with H atoms were constrained using the SHAKE algorithm. The CHARMM 36m [58] and c36 [59] force fields were applied for proteins and nucleic acids, respectively. The CHARMM modified TIP3P model [60] was used for explicit water molecules. We applied mass repartitioning to the force fields by increasing the masses of H atoms to 3 a.m.u and decreasing the masses of heavy atoms attached to H accordingly, as suggested earlier [61]. This modification allows us to use a 4 fs time step for the production run. We saved the simulation trajectories every 100 ps and obtained 5,000 frames for each replicate for +DSIF and -DSIF systems and 1,500 frames for the -DSIF_5flm system. Simulations were performed using the OpenMM program [62] on GPU machines (single NVIDIA GPU for each simulation) without any bias, restraint, or enhanced sampling using Python scripts derived from the CHARMM-GUI server [63,64].

### 2.3. Analysis of the Simulations

Simulations were analyzed for the last 400 ns (last 150 ns for -DSIF_5flm) of the production runs, since root mean square deviations (RMSD) for proteins and nucleic acids were relatively more stable after the first 100 ns of the simulations (Figs. S1-2). We analyzed the root mean square fluctuations (RMSF), contact and distance maps, pairwise distance distributions, radius of gyration (R_g_), free energies for distances and conformations, bending angles for the BH domain, Watson-Crick hydrogen bonds, probability distributions of base stacking distance and nucleic acid dihedral angles and dynamic cross-correlation matrix (DCCM) for pairs of protein domains and nucleic acids. For nucleic acids, we analyzed upstream DNA, residues from -10 to -23 and nascent RNA, the first 12 residues from the 5’ end (see Fig. 1). The first 12 residues of RNA were numbered as 1-12 in 5FLM and 31-42 in 5OIK structures.

**Figure 1.**
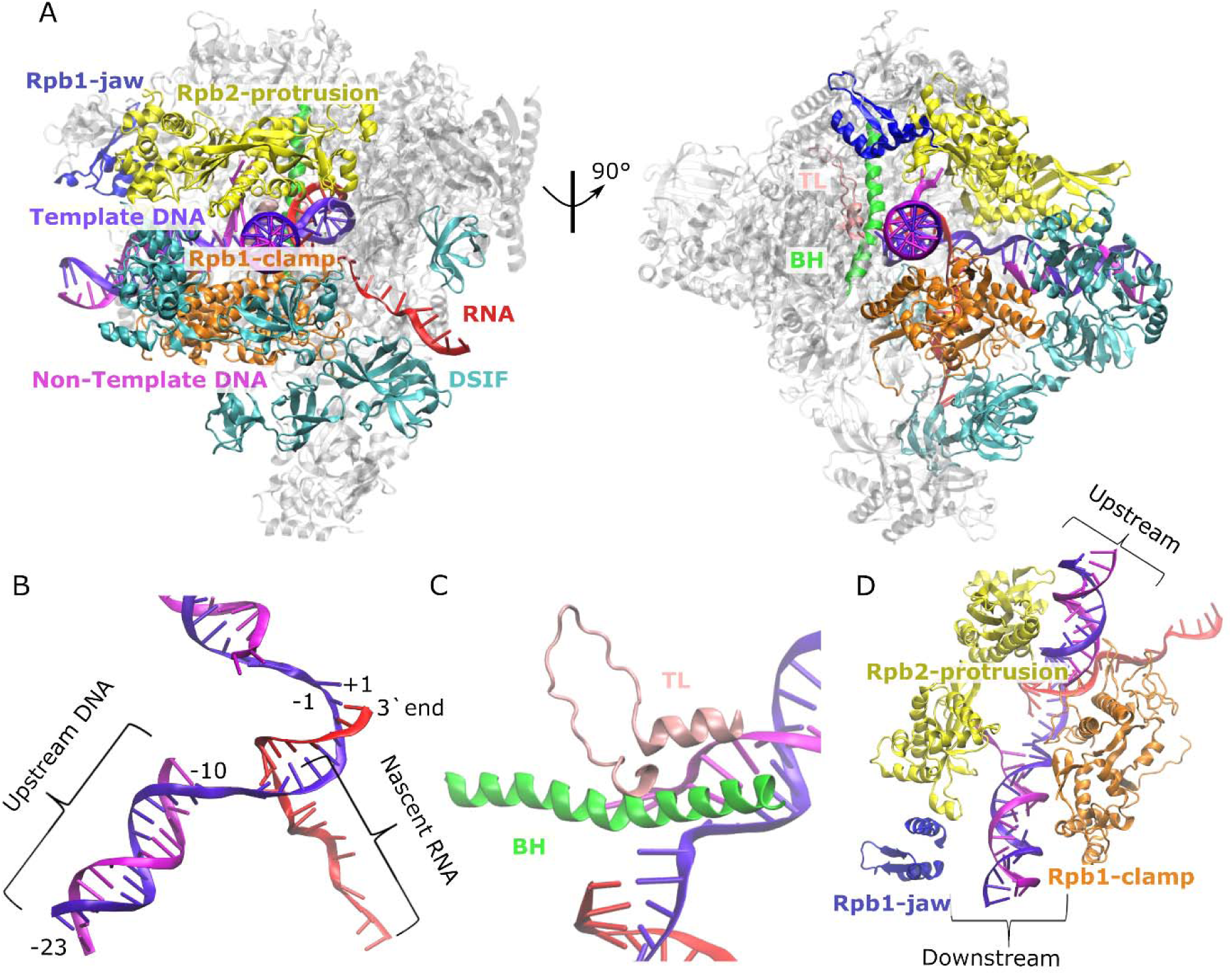
Structure of the initial Pol II – DSIF system. Structure of the complete complex (PDB ID: 5OIK) with DSIF, DNA, RNA, Rpb1 clamp and jaw domains and Rpb2 protrusion domain highlighted (A). The focused regions of the complex were zoomed in, which are upstream DNA and nascent RNA shown within curly parenthesis (B), TL and BH domains at the active site (C) and Rpb1-clamp, Rpb1-jaw and Rpb2-protrusion domains (D).

RMSD values were calculated for C_α_ and P atoms after superimposing them to the initial structures for proteins and nucleic acids, respectively. RMSF values were calculated for the same atoms with RMSD, but they were superimposed against the average structure throughout the trajectory.

Distance maps and contact maps were calculated over the trajectories and averaged over the replicates for each system. For the contact maps, two residues were assigned to have contact in a specific time frame if the minimum distance between heavy atoms of the residues is within 5 Å.

To obtain free energy for conformations of TL, BH, upstream DNA and nascent RNA, we first extracted the coordinates of C_α_ and P atoms of each amino acid or nucleotide, respectively, over the trajectory after superimposing them to the initial structure. Then, we applied principal component analysis (PCA) to reduce the dimensionality. We considered the first and second principal components (PCs) for free energy calculation. We calculated free energies by applying the weighted histogram analysis method (WHAM) using the WHAM package [65]. For the free energy based on Pol II clamp distances, we calculated the minimum distances between heavy atoms for each pair over the trajectory and then applied PCA to reduce dimensionality. This analysis was applied for two sets of distances: one is the distances between Rpb1 clamp residues from 280 to 315 and Rpb2 protrusion residues from 405 to 445 and the other is the distances between Rpb1 clamp residues from 118 to 155 and Rpb1 jaw residues from 1229 to 1297 (except the missing residues of the jaw domain in the structure).

We calculated the bending angle of BH as the angle between the principal axes of helices with residue selection of 835-852 and 853-870. For this, we first determined the covariance matrix of the C_α_ coordinates of the residues within each helix and then calculated eigenvalues and eigenvectors of the covariance matrix. After doing this calculation for the two helices, we calculated the angle between the first principal vectors, the eigenvectors with the highest eigenvalues for each helix, for the two helices from their dot products.

Watson-Crick hydrogen bonds were calculated for the upstream DNA. Hydrogen bonds were calculated between donor and acceptor atoms when they are within 1.2 Å and with a cutoff of 150° for the angle between acceptor-hydrogen-donor atoms. Base-staking distances were calculated between the neighboring bases as their center of mass distances for the heavy atoms. The dihedral angles were calculated for each nucleotide for both non-template and template DNA and distributions were calculated over the trajectory.

DCCMs were calculated for C_α_ and P atoms for proteins and nucleic acids, respectively, from the normalized covariance *C(i,j)* = < Δ**r**_i_ Δ**r**_j_ > / (< Δ**r** ^2^ > < Δ**r** ^2^ >)^1/2^, where **r**_i/j_ is the displacement vector from the average position and angle brackets denote the average over the trajectory frames. Correlated motion changed between -1 to 1, which means that the motions of two residues are negatively or positively correlated, respectively.

Analyses were performed using the Python programming language together with Scikit-learn [66], NumPy [67], and MDAnalysis [68,69] Python libraries, and the MMTSB package [54]. The VMD program [70] was used to visualize the structures obtained from the simulations.

## 3. RESULTS

In this study, we investigated the impact of DSIF on the dynamics and structure of the Pol II elongation complex to gain mechanistic insights into the role of DSIF in transcription processivity. We performed MD simulations in the presence and absence of DSIF and probed the effects of DSIF on three impact points (Fig. 1), which are the upstream nucleic acids, Rpb1-clamp, and active sites domains.

### 3.1. DSIF stabilizes DNA and RNA at exit sites

Upstream DNA and nascent RNA are known to interact with DSIF, as determined by the cryo-EM studies [12,22,23,28]. At the DNA exit, DSIF forms a “DNA clamp” by Spt5-NGN and KOW1 domains and Spt4 together with Rpb1-wall and Rpb2-protrusion domains [22,23]. Similarly, at the RNA exit, the KOW4 and KOW5 domains form an “RNA clamp” presumably to help the proper exit of RNA [22,23]. We performed analyses to investigate how the interactions with DSIF impact the dynamics of DNA and RNA. RMSF analyses suggested that DNA (template and non-template) has reduced RMSF in the presence of DSIF (Fig. 2A), suggesting a reduced flexibility for the DNA at the exit site. However, differences in the RMSF value of RNA are less significant since there are large error bars, suggesting that fluctuations differ for each replicate simulation. The impact on the RMSF is stronger for DNA, which could be partly due to the overall higher dynamics of the single-stranded nascent RNA compared to base-paired DNA. The increased stability suggested by RMSF values could be the result of the impact of the increased interactions between protein and nucleic acids. The total number of contacts for DNA increased more than twofold for +DSIF, in which DSIF makes 74 % of the contacts with the upstream DNA (Table 1) and those contacts are mostly with Spt4, Spt5-NGN and KOW1 domains (Fig. S3). On the contrary, for RNA, the total number of contacts slightly decreased for +DSIF (Table 1). Spt5 started to have a large percentage of contacts, mostly with the KOW4 domain (Fig. S4), that possibly play a role in stabilizing RNA at the exit site. On the other hand, the contacts with the Rpb2-wall domain were significantly reduced in the presence of DSIF (Fig. S4B).

**Figure 2.**
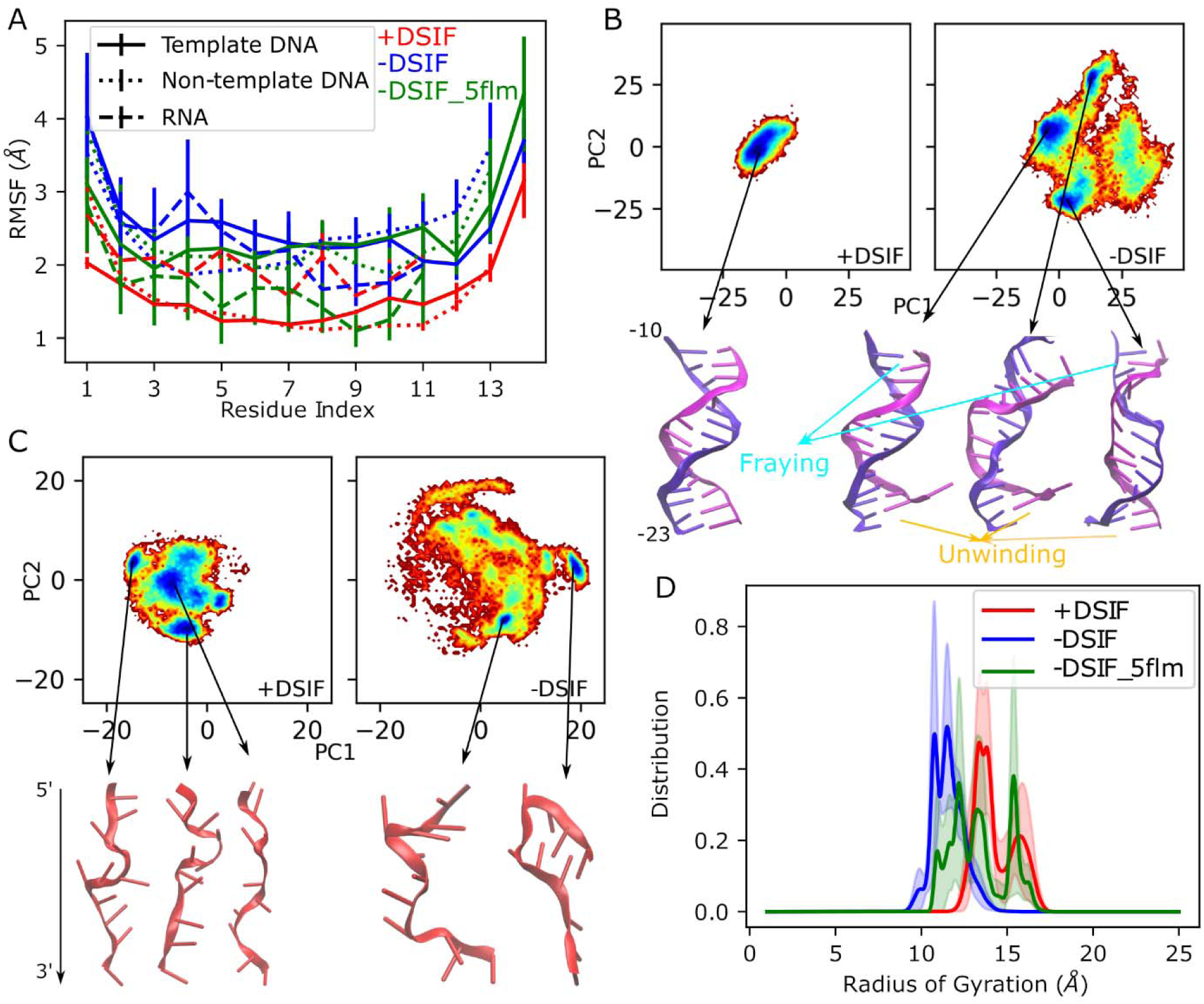
Structure and dynamics of upstream DNA and nascent RNA. RMSF values were reduced for DNA in the presence of DSIF (A). Free energy profiles of DNA (B) and RNA (C) conformations showed more stable structures for DNA and more extended structures for RNA in the presence of DSIF. Free energy profiles were calculated from the first two principles components (PC1 and PC2) from the PCA based on Cartesian coordinates (see Methods for details). The radius of gyration analysis for RNA (D) suggests increased number of compact structures in the absence of DSIF.

**Table 1.** Number and percentage of contacts of DNA and RNA with protein.

| +DSIF |  |  |  |  |
| --- | --- | --- | --- | --- |
| Protein | Number of DNA contacts | Percentage of DNA contacts | Number of RNA contacts | Percentage of RNA contacts |
| Total | 109.98 ± 5.68 | 100 | 112.99 ± 3.99 | 100 |
| Spt4 | 18.94 ± 3.60 | 17.22 | 0.0 ± 0.0 | 0 |
| Spt5 | 62.36 ± 5.32 | 56.70 | 35.05 ± 3.79 | 32.02 |
| Rpb1 | 5.91 ± 1.93 | 5.38 | 28.92 ± 4.77 | 25.60 |
| Rpb2 | 22.76 ± 1.37 | 20.7 | 49.02 ± 2.69 | 43.38 |
| -DSIF |  |  |  |  |
| Protein | Number of DNA contacts | Percentage of DNA contacts | Number of RNA contacts | Percentage of RNA contacts |
| Total | 46.90 ± 7.38 | 100 | 119.09 ± 6.26 | 100 |
| Rpb1 | 12.53 ± 8.14 | 26.72 | 50.29 ± 4.71 | 42.23 |
| Rpb2 | 34.37 ± 6.05 | 73.28 | 68.77 ± 3.95 | 57.75 |
| -DSIF_5flm |  |  |  |  |
| Protein | Number of DNA contacts | Percentage of DNA contacts | Number of RNA contacts | Percentage of RNA contacts |
| Total | 38.88 ± 6.97 | 100 | 123.22 ± 11.79 | 100 |
| Rpb1 | 12.09 ± 7.85 | 31.10 | 60.78 ± 5.46 | 49.33 |
| Rpb2 | 26.78 ± 5.89 | 68.90 | 61.34 ± 11.50 | 49.78 |

The number of contacts was calculated over the trajectories and averaged over the number of replicate simulations. DNA contacts were calculated for the upstream DNA (from -10 to -23). RNA contacts were calculated for the first 12 residues of nascent RNA where it contacts with DSIF.

Next, we analyzed the structural changes occurring in DNA and RNA in the presence of DSIF. We observed larger RMSD values in the absence of DSIF (Fig. S2), which suggests that nucleic acid structures are deviating from the initial structure in the absence of DSIF. To understand the nature of structural deviation, we analyzed the conformations of the upstream DNA and nascent RNA. We performed PCA on the Cartesian coordinates of P atoms of DNA and RNA from the superimposed frames along the trajectory. Fig. 2B shows that the DNA conformation is mostly stable in the presence of DSIF. On the other hand, a broader conformational space was observed for DNA when DSIF was removed. The minimum energy structure for the +DSIF system shows a base-paired DNA structure without any significant fraying or unwinding. However, the three minimum energy structures for the -DSIF system show significant deviation from the initial structure and fraying and unwinding were observed. We observed a similar conformational deviation for the DSIF_5flm system, for which two populated structures showed significant unwinding (Fig. S5A). To better understand the structural changes, we analyzed Watson-Crick base-pair hydrogen bonds, base stacking distances, and dihedral angles. The number of base-pair hydrogen bonds slightly decreased for the -DSIF system and significantly reduced for the -DSIF_5flm system (Fig. S6A). For the -DSIF_5flm system, large errors suggest that some of the replicates showed significantly decreased base-pairs than the others, which could be a result of an unstable initial structure due to the modeling of upstream DNA for this system. We also observed a partial increase in base stacking distances, which appears as a small shoulder in the distance distribution plot of -DSIF and -DSIF_5flm systems (Fig. S6B). These increased base stacking distances reflect the fraying we observed in the most populated structures shown in Figs. 2B and S5A. For the dihedral angles, we observed changes in zeta and chi angles, which show slightly broader distributions for the -DSIF and -DSIF_5flm systems (Fig. S7). The changes in these angles further suggest a deviation from the base-pair structures in the absence of DSIF.

RNA conformations were sampled in large spaces for both systems, although for -DSIF, the conformational space was somewhat broader (Fig. 2C). The minimum energy structures for +DSIF are mostly elongated, while for the -DSIF system, the structures are more compact, with one structure forming a hairpin. Similar results were observed for -DSIF_5flm, such that the mostly populated structure was compact and hairpin formation was observed for the second populated region (Fig. S5B). To understand the impact of DSIF on the compactness of RNA structure, we calculated R_g_ values over the simulation trajectories. We observed that R_g_ values are significantly lower for the -DSIF system and lower Rg values were also observed for the -DSIF_5flm system (Fig. 2D), suggesting that DSIF prevents RNA from forming compact structures, including hairpins.

Overall, our results suggest that DSIF forms strong interactions with upstream DNA and nascent RNA, reduces the dynamics of DNA at the exit tunnel, and prevents them from forming unusual structures, like frayed base-pairs for DNA and hairpin-like secondary structures for RNA.

### 3.2. DSIF alters the dynamics and interactions of TL and BH

DSIF does not have direct interactions with TL and BH, both are near-active site domains. However, earlier studies suggested that DISF may affect the Pol II active site allosterically [6,24,31,33,37], though the molecular basis of such allosteric impacts is not known. To explore the impacts of DSIF on the active site, we analyzed the structure and dynamics of TL and BH. RMSF plots showed increased flexibility for TL in the presence of DSIF especially when comparing with the -DSIF system, without any significant change for BH (Fig. 3A). The increased flexibility for TL was observed in the loop region between residues 1109-1115, which was shown in color in Fig. 3B. TL is known to play a key role in the nucleotide addition cycle (NAC), and it alternates between “open” and “closed” structures during the NAC [40,44,71–77]. TL in the closed state is proposed to facilitate nucleotide selection and position the incoming nucleotide for catalysis, while the open TL is proposed to play roles in nucleotide binding and forward translocation. In the cryo-EM structure used in our simulations, TL is partially unresolved due to high flexibility in its open state. We modeled the unresolved part and showed the comparison with the closed state in Fig. 3C. In the presence of DSIF, increased flexibility occurred in this modeled region, which is the tip of the TL that normally moves toward to active site to form the closed state. Based on the increased dynamics on the tip of TL in the presence of DSIF, we hypothesize that DSIF might help facilitate the open-closed TL transition, potentially stabilizing the closed TL.

**Figure 3.**
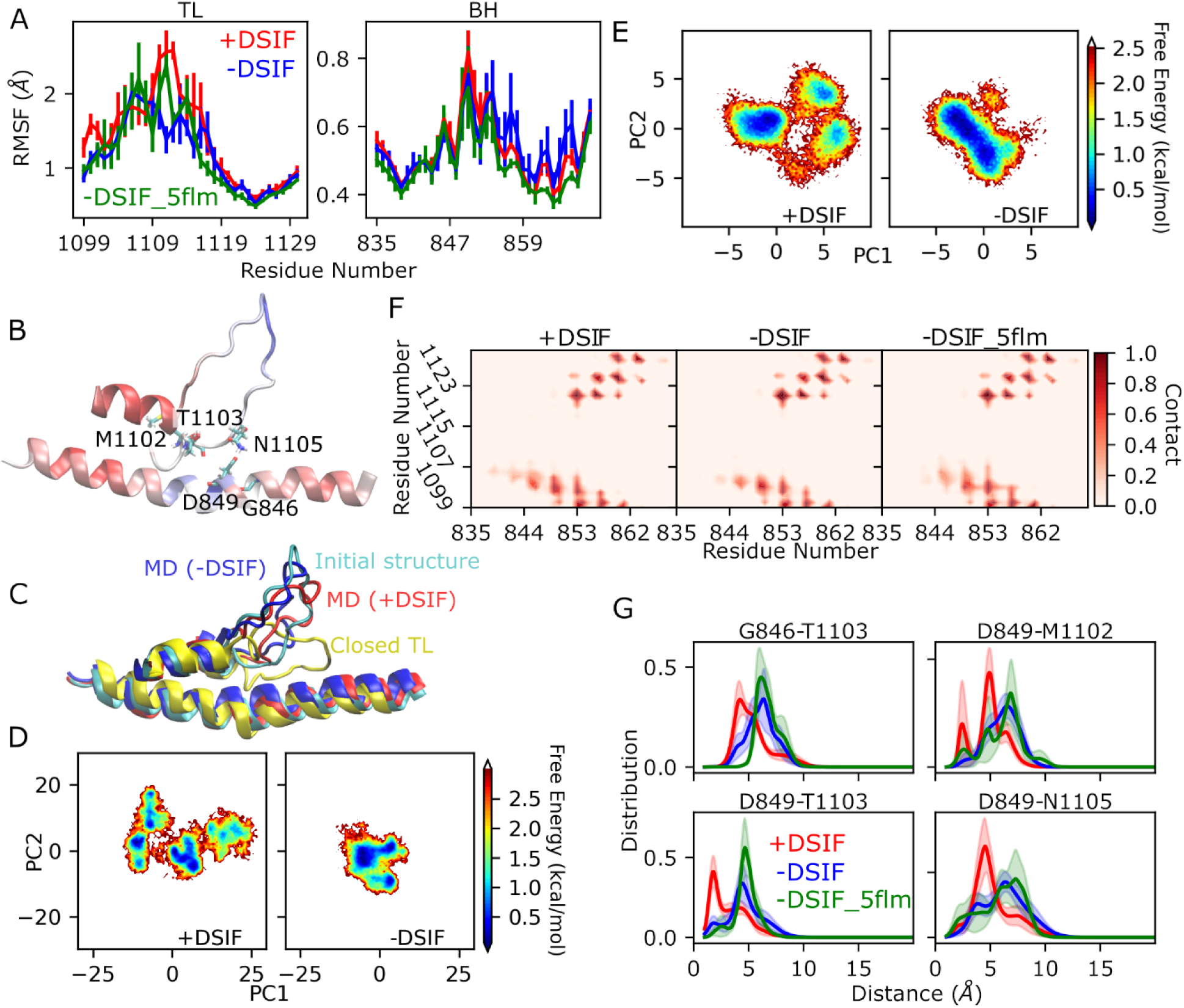
Structure and dynamics of TL and BH. RMSF values for BH and TL showed differences only for TL with increased flexibility in the presence of DSIF (A), and RMSF values are shown in colors from red to blue in increasing order in the structure of TL and BH, the color ranges were RMSF values of 0.40-0.82 for BH and 0.60-2.58 for TL (B). The initial modeled structure of TL, the closed TL structure from PDB ID: 2E2H, and the structures from +DSIF and -DSIF simulations extracted based on PCA are superimposed (C). PCA showed larger conformational spaces for both TL (D) and BH (E), and the structure for TL showed larger deviation from the initial model in the presence of DSIF (C). The number of contacts between BH and TL increased in the presence of DSIF (F), and in consistency with the number of contacts, the distances between selected TL-BH pairs were reduced (G). These selected pairs are located at the BH-TL interface that have high flexibilities, which are colored by RMSF values from the +DSIF system (B).

To gain more insight into the effects of DSIF on TL conformation, we performed PCA on TL C_α_ coordinates. Fig. 3D shows that TL samples a larger conformational space in the presence of DSIF, which is consistent with higher dynamics observed in the RMSF plots. The free energy plots show multiple minimum energies. We extracted the structures of TL that are closest to the globular minimum in the free energy plots. We observed that for the +DSIF system, the minimum energy structure showed a larger deviation from the initial modeled structure (Fig. 3C), as expected from the larger deviations in the RMSD plots (Fig. S1). BH also sampled a somewhat larger conformational space for the +DSIF system (Fig. 3E), while the lowest energy appears in the same region for both systems, and the corresponding conformations extracted from the trajectories are also structurally similar, with bending angles of 156.2° and 159.7° for +DSIF and -DSIF systems, respectively. In addition, bending angle distributions throughout the trajectories are broad with large errors and, therefore, did not show any significant differences for +DSIF and -DSIF systems (Fig. S8). On the other hand, the -DSIF_5flm system showed sharper distribution for the BH bending angles and the maximum angle (161.2°) appears close to the maximum angle of -DSIF (160.9°) and smaller compared to +DSIF (164.0°). This suggests that in the absence of DSIF, BH tend to bend more, while in the presence of DSIF, BH samples between bent and straight conformations manifested themselves as a broad bending angle distribution.

Although we did not observe any significant changes in the BH structure and dynamics, we observed changes in its interactions with TL. The number of contacts between the two domains increased for the BH residues between 840-849 and the TL residues between 1102-1105 for +DSIF system (Fig. 3F). To understand which contacts significantly differ for the two systems, we calculated the distance distributions for the pairs that have normalized number of contacts more than 0.7 in any of the two systems and the difference of the contacts between the two systems is larger than 0.1. Fig. 3G shows the distance distributions for the pairs that show the most significant changes for the two systems, as well as for the -DSIF_5flm system, while the distances for the other pairs were only moderately different (Fig. S9). We observed that BH and TL distances are similar for -DSIF and -DSIF_5flm systems and decreased for the +DSIF system, suggesting increased interactions between the two domains for the TL residues M1102, T1103 and N1105 and the BH residues G846 and D849 in the presence of DSIF. The positions of these residues were shown in Fig. 3B. An earlier MD simulation study suggested that TL-BH interactions around that region of the TL play an important role in the TL dynamics [45] supporting our hypothesis that DSIF may play a role in the TL opening/closing motion. Overall, our results suggest that DSIF may increase the dynamics of TL and its interactions with BH and decrease the bending of BH.

### 3.3. DSIF regulates the clamp conformation at the central cleft

Earlier studies suggest that DSIF may help maintain the clamp in its closed conformation, thereby enclosing the DNA for transcription elongation [24,34,35,78]. To understand the impact of DSIF on the size of the central cleft, we analyzed the distances of the Rpb1-clamp to Rpb1-jaw and Rbp2-protrusion at downstream and upstream, respectively. We calculated all the pairwise distances for the selected helical regions of the clamp and jaw and the coil-coiled domains of the clamp and protrusion (Fig. 4A, the residue selection is provided in the Methods section). Fig. 4B shows the free energy profiles of the PCs that were obtained by applying PCA on the pairwise distances. Energy profiles span similarly large spaces for +DSIF and -DSIF systems for clamp-jaw distances, while for clamp-protrusion distances, the +DSIF system spans a narrower space, suggesting that direct interactions of DSIF with the clamp stabilize clamp motion at the upstream.

**Figure 4.**
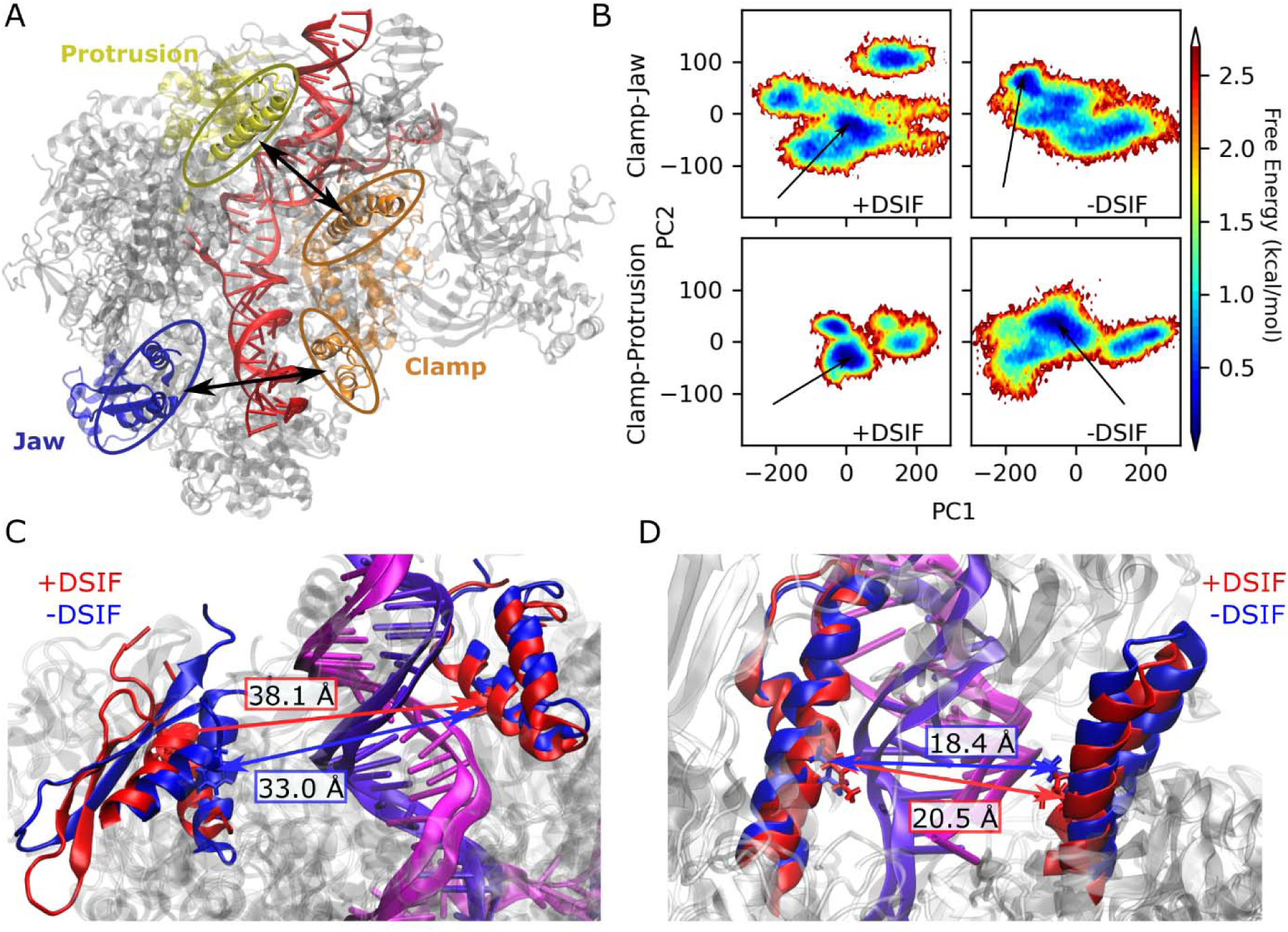
Analysis of clamp motion. The architecture of the central cleft formed by Rpb1-clamp and other domains is demonstrated in a cartoon representation (A). The free energy profiles of PCA based on pairwise distances between Rpb1-clamp and Rpb1-jaw, and between Rpb1-clamp and Rpb2-protrusion (as colored in panel A) are shown (B). The minimum energy structures from the PCA showed that the distances increased in the presence of DSIF, both for clamp-jaw (C) and clamp-protrusion (D).

We then extracted the most populated structures based on the distance PCA free energy profiles and visualized them after superimposing the Rpb1 subunits for +DSIF and -DSIF systems (Fig. 4C-D). Clamp and jaw domains became relatively closer after we removed DSIF (Fig. 4C), suggesting that DSIF may play a role in regulating the distance between the two domains to maintain the proper size in the cleft. Consistently, distance maps also show a decrease in distances between the clamp and jaw domains in the absence of DSIF (Fig. S10). In addition, distance distribution profiles for the selected residue pairs showed significant differences between the two systems, with the +DSIF system showing overall larger distances (Fig. S11). On the other hand, -DSIF_5flm system showed somewhat increased distances in the distance map (Fig. S10) and distance profiles for selected residues show broader distributions than the other two systems, with a smaller shoulder on the larger distance region (Fig. S11). This may be due to the shorter simulation times for the -DSIF_5flm system, however, overall, the discrepancies between -DSIF and -DSIF_5flm systems may suggest that the globular conformational changes, like clamp motion, may not be fully accessible with nanosecond time scale simulations.

At the upstream, the protrusion domain slightly came closer to the coil-coiled domain of the clamp in the absence of DSIF (Fig. 4D). Consistent with the extracted structures, distance maps also show reduced distances for both -DSIF and -DSIF_5flm (Fig. S10). More prominently, we observed sharper distance profiles (Fig. S12) in the presence of DSIF compared to both -DSIF and -DSIF_5flm systems, suggesting that DSIF stabilizes the coil-coiled domain of the clamp. Since there are direct interactions between DSIF and the coil-coiled domain (Fig. S13), we concluded that direct interactions reduced the dynamics of the coil-coiled domain and consequently caused sharper distance profiles. Overall, our results suggest that DSIF may help maintain the distances of the clamp to the other side of the cleft, while removing DSIF decreases these distances, which may potentially impact transcription processivity. In addition, the initial structures with and without DSIF have similar clamp conformations (Table S1), and the pairwise distances of clamp residues to the jaw and protrusion residues were similar for the two structures (cyan and purple lines in Figs S11-12). In our simulations, we observed overall reduced pairwise distances compared to initial structures with a broader distribution for the systems without DSIF. Therefore, the simulations provided a range of dynamics for clamp motion and suggested that DSIF partly reduces those dynamics. On the other hand, we observed discrepancies in the distances between the clamp and jaw domains for the simulations of the two systems without DSIF. Therefore, we acknowledge that our simulations may not provide results relevant to long time scale globular motions including clamp motion at the central cleft.

### 3.4. Motions of active site loops are correlated with clamp and RNA motions

The impacts of DSIF on the active site have been associated with the allosteric effects of either nucleic acids or Rpb1-clamp. To understand the presence of any allosteric effect, we calculated the correlated motions (see Methods for DCCM calculations) for TL and BH with upstream nucleic acids and Rpb1-clamp. We provided DCCM maps colored in red for both positive and negative correlations in Fig. 5A, while the colors coded by the sign of correlations can be seen in the supplementary figures (see Figs. S14-16). First, we calculated the correlated motion between BH and TL and observed that DSIF increased the correlation between them compared to -DSIF and -DSIF_5flm systems (Figs. 5A and S14), consistent with the increased interactions between these two domains. For the nascent RNA, in the absence of DSIF, for both -DSIF and -DSIF_5flm systems, correlations with TL and BH increased (Figs 5A, S15 and S16), especially for the regions shown with the green arrow in Fig. 5A. Such an increase in the TL/BH-RNA correlated motions could be related to RNA hairpin formation as studies suggested that hairpin formation in the exiting RNA may have an increased effect on the active site structure and dynamics [79–85]. We also observed somewhat increased correlated motions between DNA and TL (Fig. S16), which suggests that both DNA and RNA may have an allosteric impact on TL motion.

**Figure 5.**
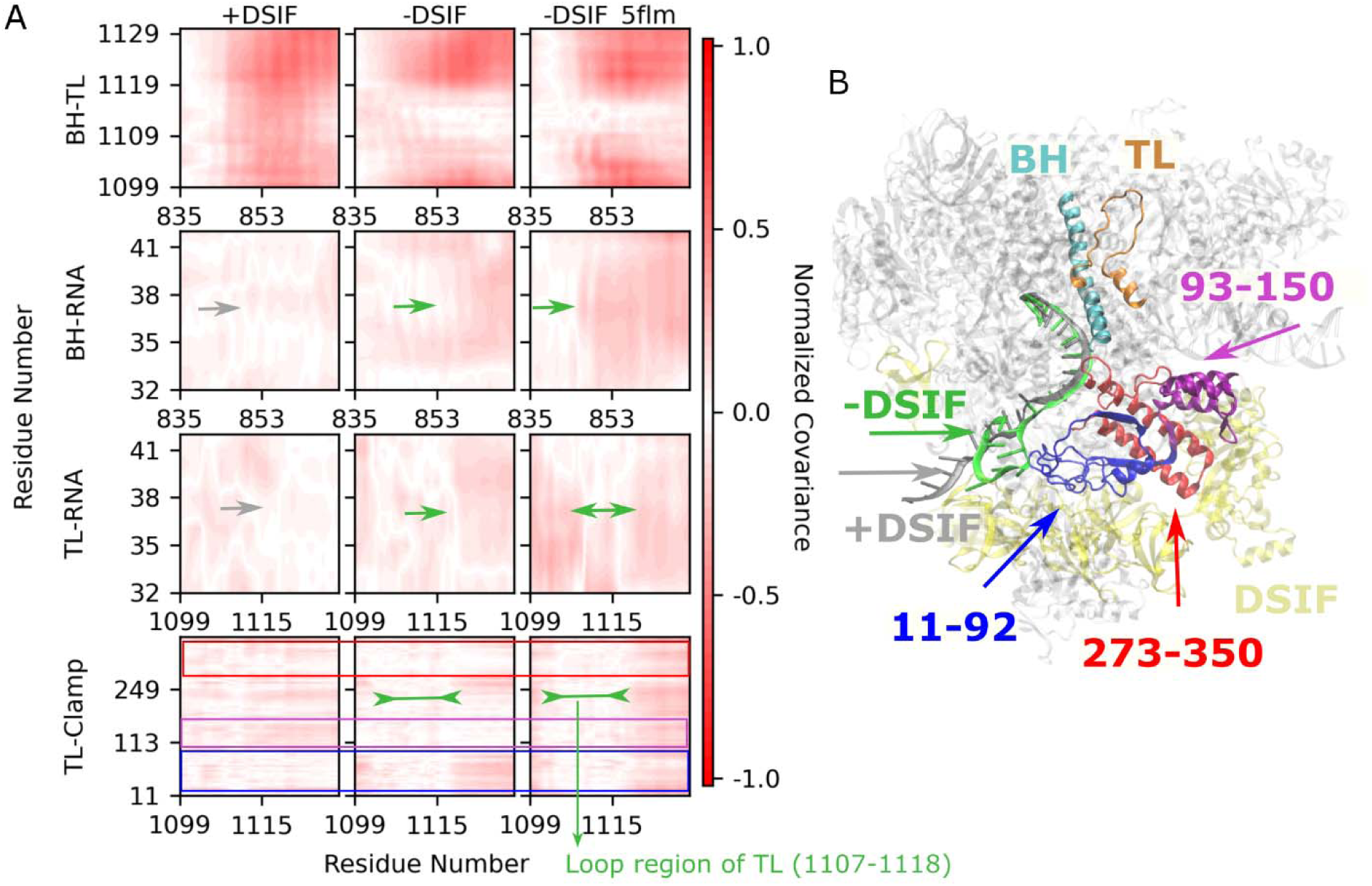
DCCM analysis for protein domains and nucleic acids. Correlated motions decreased between BH and TL and increased between BH and RNA, and TL and RNA in the absence of DSIF (A). In the presence of DSIF, correlated motion decreased for TL with clamp residues 11-92, increased with clamp residues 93-150 and 273-350, with the correlations of the TL loop region was most impacted (A). Secondary structure formation of RNA and motion of clamp residues correlated with the TL dynamics, suggesting an allosteric impact of DSIF on TL motion (B). The residue ranges 11-92, 93-150, and 273-350 in panel B are highlighted in the same color as cross correlation plot for TL and clamp in panel A to emphasize that these are the regions with changes in cross-correlation maps.

For the clamp, we observed changes in three regions in the correlation motions with and without DSIF for TL (Fig. 5A) and similarly for BH (Fig. S14). DSIF decreased the correlations with TL for the clamp residues 11-92. This region interacted with KOW2-4 domains of the DSIF (Fig. S17), and these interactions potentially reduced its dynamics (Fig. S18). Therefore, we concluded that decreased dynamics reduced the impact of this region on the active site motion. On the contrary, the regions with residues 93-150 and 273-350, which include the coiled-coil domain and close-by residues, showed increased correlations with TL in the presence of DSIF (Figs. 5A and S14). More strikingly, we observed decreased correlations between the clamp and TL loop region (residues 1107-1118) for both -DSIF and -DSIF_5flm systems. This suggests that DSIF impacts the loop region of the TL allosterically by altering its correlated motion with the Rpb1 clamp.

Overall, we suggest that the collective motion of the clamp residues and RNA structural changes impacts motions of TL and BH in the presence of DSIF (Fig. 5B). DSIF affected the correlations of the TL with the clamp residues and, potentially, these correlated motions caused increased flexibility of the TL that we observed in the RMSF plots (Fig. 3A). In addition, DSIF prevents RNA from forming hairpin-like secondary structures and, by this, reduces the impact of RNA on the active site dynamics.

## 4. DISCUSSION

DSIF is the only universally conserved elongation factor, which is known to have regulatory functions in various steps of transcription. Although it is widely studied, its regulatory mechanism remains unresolved. To obtain mechanistic insights into the DSIF function at the molecular level, we performed MD simulations of the Pol II elongation complex in the presence and absence of DSIF. The most notable impact of DSIF was observed in the nucleic acid structure and dynamics at the exit sites. We observed that DSIF stabilized the upstream DNA, causing it to sample a narrower conformational space, preventing it from forming frayed states and potentially unwinding. DSIF’s impact on maintaining upstream DNA structure may be a contributing factor to its role in processivity. Structural studies suggest that DSIF forms a DNA clamp at the upstream together with Rpb1-wall and Rpb2-protrusion domains [22,23]. Forming the DNA clamp helps stabilize DNA at the exit site, proposed to prevent it from unwinding and potentially rewinding with exiting RNA. Our observation of reduced unwinding of the DNA base pair at the exit site in the presence of DSIF supports this earlier hypothesis on the role of the DNA clamp formed by DSIF. In addition, DSIF demonstrated an even larger impact on the structure of the nascent RNA, preventing it from forming more compact secondary structures, including hairpins. Earlier studies suggest that the hairpin formation of RNA allosterically affects the polymerase active site and stimulates pausing [79–85]. In the light of these earlier studies, our results suggest that DSIF may promote processivity by helping RNA maintain more extended secondary structures at the exit site and consequently preventing RNA from allosterically causing the formation of paused states.

DSIF has also been proposed to impact the clamp motion, potentially stabilizing a closed clamp and, in this way, maintaining processivity [24,34,35,78]. Therefore, we expected to observe the opening of the clamp in our simulations after removing DSIF. On the contrary, we observed a slightly more closed clamp as the distances between the clamp and the domains on the other side of the cleft reduced in the absence of DSIF. Additionally, the clamp, especially the CC domain of the clamp, becomes more dynamic in the absence of DSIF, suggesting that DSIF may play a role in stabilizing the CC domain and, consequently, clamp motion. Based on our results, we propose that, instead of further closing the clamp, DSIF may help maintain the proper size for the central cleft by stabilizing the clamp motion and preventing it from opening or closing further to form unfunctional conformational states. However, we also note that our conclusion is limited to the simulation time scale, which may not be sufficient to completely sample the globular conformational changes in the central cleft. Especially, the discrepancies in the clamp-jaw distances from the simulations of -DSIF and -DSIF_5flm systems suggest that the clamp motion may not be accurately captured for the nanosecond simulation time scale.

Furthermore, we observed changes in the dynamics of TL and the interactions between TL and BH. TL showed increased flexibility for the residues from Y1109 to K1115 (F1086 to K1092 in yeast), in the presence of DSIF. Previous studies suggest that residues F1086 and V1089 of the TL in yeast Pol II demonstrated loss-of-function mutations [86], and earlier structures [87] suggest an important role for these residues in TL closing dynamics as partial closing of the TL correlated with the loss of the hydrogen bond between the backbones of F1086 and V1089 (Y1109 and V1112 in the mammalian structure). We did not observe any hydrogen bond formation for the backbones of the two residues in either of the simulations. However, the increased flexibility of the TL tip, including the residues Y1109 and V1112 in the presence of DSIF, may suggest DSIF’s impact on the TL closing motion. We also observed increased interactions between BH and TL, especially for TL residues M1102, T1103, and N1105 and BH residues G846 and D849. BH and TL interactions are known to be important for the catalytic function [38,76,77,86] and transition between closed and open TL [45]. Also, a decrease in BH-TL interactions was proposed to be one of the mechanisms for the loss-of-function phenotypes of TL mutants [38]. Here, based on our results, we hypothesize that DSIF impacts the TL opening/closing dynamics, which is not feasible to model by standard MD simulations that we applied here. One of our future directions is to perform enhanced sampling simulations to investigate the impacts of DSIF on the energetics of the transition between the open and closed TL, as has been done in an earlier study performed in the absence of DSIF [45].

DSIF was proposed to affect the translocation dynamics during the nucleotide addition cycle [6,25,36,88], Studies suggested that NusG, the bacterial homolog of Spt5, stabilized the post-translocation state by presumably affecting the active site domains [33,37]. We observed three main impacts of DSIF on active site domains, which are increased loop dynamics of the TL, increased interactions between TL and BH, and broader bending angle distributions of BH. The first two effects on TL suggest that there is a destabilizing effect on TL in its open state, which would be accompanied by increased TL motion toward BH and formation of BH-TL contacts. According to the proposed nucleotide addition cycle, Pol II oscillates between pre-and post-translocation states when the TL is open and the concerted motion of TL and BH moves Pol II to the post-translocation state [39,89]. Our observation of increased TL dynamics and TL/BH interactions may indicate a change in the pre-post translocation equilibrium towards post-translocation and, therefore, may support the earlier hypothesis on stabilization of post-translocation by DSIF. On the other hand, BH bending is proposed to be coupled with TL closing to support post-translocation [37,90]. Our results suggest a broader bending profile compared to the systems without DSIF, with overall less bending. This is not consistent with the proposed role of the BH bending in stabilizing the post-translocation state. However, increased dynamics on the BH bending and straightening may also play a role in affecting the equilibrium between pre-and post-translocation states by the impact of DSIF, as such a role for NusG was suggested by previous experimental studies [33,37].

The role of DSIF in processivity has been associated with its allosteric impact on the structure and dynamics of the active site [6,24,31,33,37], especially through the Rpb1-clamp and upstream nucleic acids. To understand allosteric interactions, we calculated the correlated motions by constructing DCCM for TL and BH with the clamp and nucleic acids. Our results suggest that the clamp has altered correlated motion with TL and BH domains in the presence of DSIF. We especially observed increased correlations for the loop region of TL with the clamp residues.

This is consistent with the earlier studies that focused on the allosteric impact of the clamp and its CC domain [31,34,78]. However, the molecular mechanisms of how the clamp impacts the TL dynamics to promote processivity are not well understood. Hypotheses include altered clamp position by DSIF interactions at the coiled-coil domain affecting active site dynamics [31,91] or stabilized closed clamp conformation by DSIF altering the TL conformation to facilitate catalysis [6,80]. In addition, our results suggest that not only the clamp but also RNA structures impact TL, as in the absence of DSIF, and consequently, upon formation of hairpin-type secondary structures, cross-correlations of TL with RNA increase. Therefore, we conclude that, in the presence of DSIF, the concerted motions of DNA/RNA and Rpb1-clamp impact the active site dynamics and propose that these motions may help prevent interruptions during transcription elongation for maintaining processivity.

## 5. CONCLUSION

In this study, we examined the impacts of DSIF on the dynamics of the Pol II elongation complex by performing MD simulations in the presence and absence of DSIF. DSIF impacted the structure and dynamics of the upstream nucleic acid substantially, and Rpb1 clamp opening/closing motion moderately. We observed that motions of both nucleic acids and the clamp domain are correlated with the motion of the active site domains, TL and BH, impacting their dynamics and interaction networks. Our results suggest that DSIF may allosterically affect the TL opening/closing motion. The next step to understand DSIF’s role in the conformational interplay of TL would be to model the TL opening/closing motion using enhanced sampling techniques. Probing the effects of DSIF on the energetics of the open and closed TL and the transition pathway between the two would provide important insights into the role of DSIF in transcription processivity.

## DATA AND SOFTWARE AVAILABILITY

Software used for the simulations and analysis is all open source, which includes OpenMM, CHARMM academic version, MMTSB, and Python libraries (see Methods section for the details of the software). The structures with PDB IDs: 5OIK and 5FLM were used as the initial structures. The Python scripts to run the OpenMM program were generated using the CHARMM-GUI server. The data for ensuring reproducibility of our study is provided in the GitHub repository (https://github.com/bercemd/PolII_DSIF). The data includes coordinates of the initial solvated systems, scripts to prepare the initial systems, run input files, parameter and topology files for the simulations, analysis scripts, and the output data from the analysis. MD simulation trajectories were not included in the repository due to their large sizes; however, they can be made available to researchers upon reasonable requests.

## ACKNOWLEDGEMENTS

The authors used computational resources at the Cyberinfrastructure and Research Technologies at the University of California Merced (grant numbers: NSF ACI-2019144, NSF ACI-1429783) and at the Advanced Cyberinfrastructure Coordination Ecosystem: Services & Support (ACCESS) with ACCESS award number BIO210145.

## AUTHOR INFORMATION

### Author Contributions

W. D. A.: Research design and development, analysis, manuscript reviewing and editing; B. M. B.: Research design and development, simulations, analysis; B.D.: Research design, development and management, simulations, analysis, manuscript original writing, reviewing and editing. All authors have given approval to the final version of the manuscript.

### Note

The authors declare no competing financial interest.

## NOTES

This document is the unedited Author’s version of a Submitted Work that was subsequently accepted for publication in the *Journal of Chemical Information and Modeling*, copyright *© 2025 American Chemical Society* after peer review.

## SUPPORTING INFORMATION

Supplementary material associated with this article contains the following analysis results: RMSD for Pol II-Rpb1, TL, BH, RNA and upstream DNA; RMSD of the protein domains for the two initial PDB structures; contact maps of DNA and RNA with the nearby protein domains at the upstream and protein-protein contact maps of Rpb1-clamp with KOW2-4 and CC domain with NGN and KOW2; probability distributions of the number of Watson-Crick base-pair hydrogen bonds, base stacking distances and nucleic acid backbone dihedral angles; probability distributions of BH bending angles; probability distributions of BH-TL distances, distances of Rpb1-clamp to Rpb1-jaw and Rpb2-protrusion; distance maps between Rpb1-clamp and Rpb1-jaw, and Rpb1-clamp and Rpb2-protrusion; cross-correlation maps of TL and BH with each other, and with Rpb1 clamp, DNA and RNA; RMSF values for Rpb1-clamp.

