## Supporting Information for "Molecular Basis for Impacts of DSIF on the Dynamics of RNA Polymerase II Elongation Complex"

Supplementary material associated with this article contains the following analysis results: RMSD for Pol II-Rpb1, TL, BH, RNA and upstream DNA; RMSD of the protein domains for the two initial PDB structures; contact maps of DNA and RNA with the nearby protein domains at the upstream and protein-protein contact maps of Rpb1-clamp with KOW2-4 and CC domain with

NGN and KOW2; probability distributions of the number of Watson-Crick base-pair hydrogen bonds, base stacking distances and nucleic acid backbone dihedral angles; probability distributions of BH bending angles; probability distributions of BH-TL distances, distances of Rpb1-clamp to Rpb1-jaw and Rpb2-protrusion; distance maps between Rpb1-clamp and Rpb1-jaw, and Rpb1-clamp and Rpb2-protrusion; cross-correlation maps of TL and BH with each other, and with Rpb1 clamp, DNA and RNA; RMSF values for Rpb1-clamp.

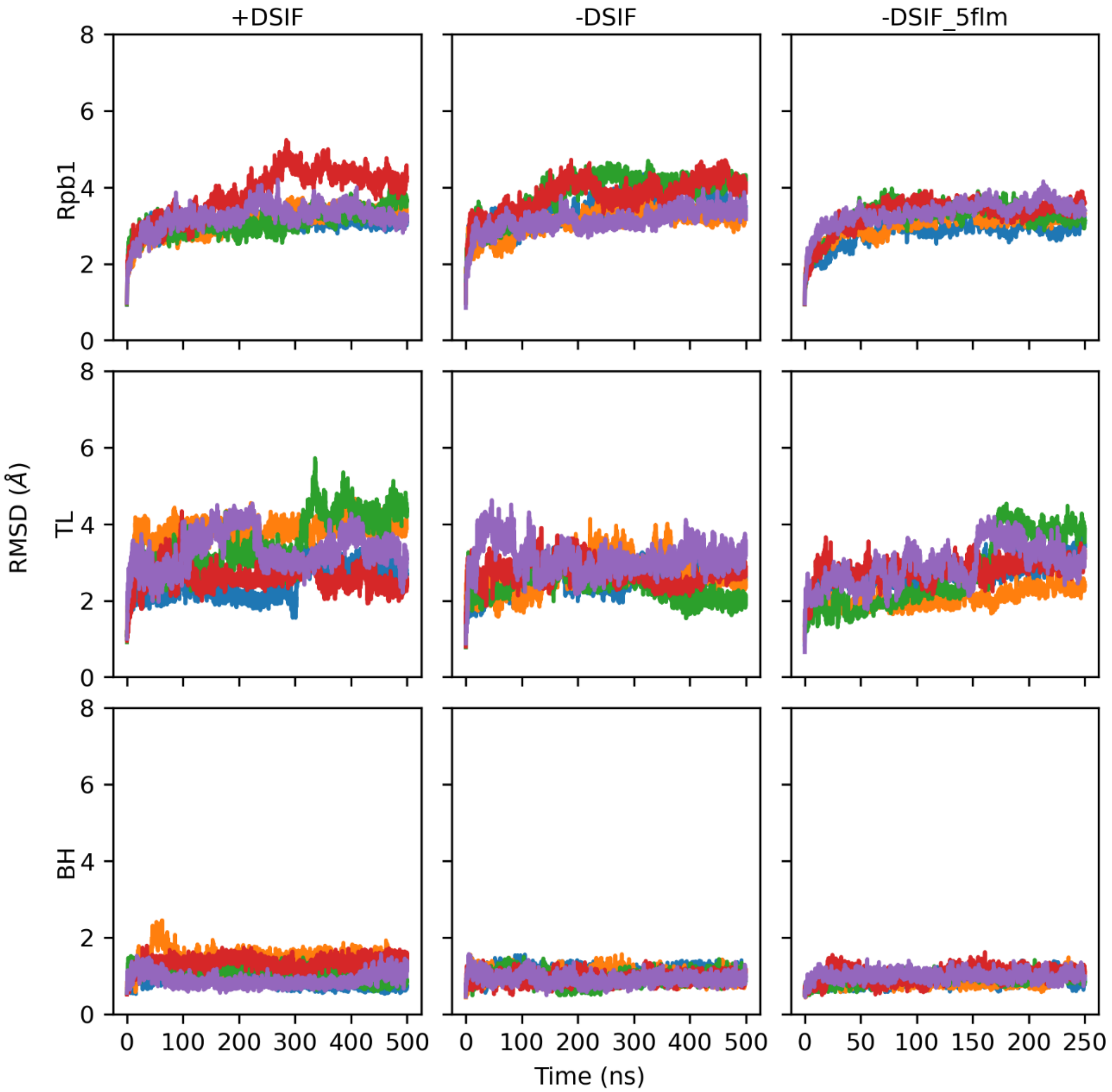


**Figure S1.** The root mean square deviation (RMSD) of RNA polymerase II (Pol II) Rpb1 subunit, TL and BH domains of Rpb1 for the systems +DSIF, -DSIF and -DSIF_5flm. Lines with different colors represent RMSD from each replicate simulation.

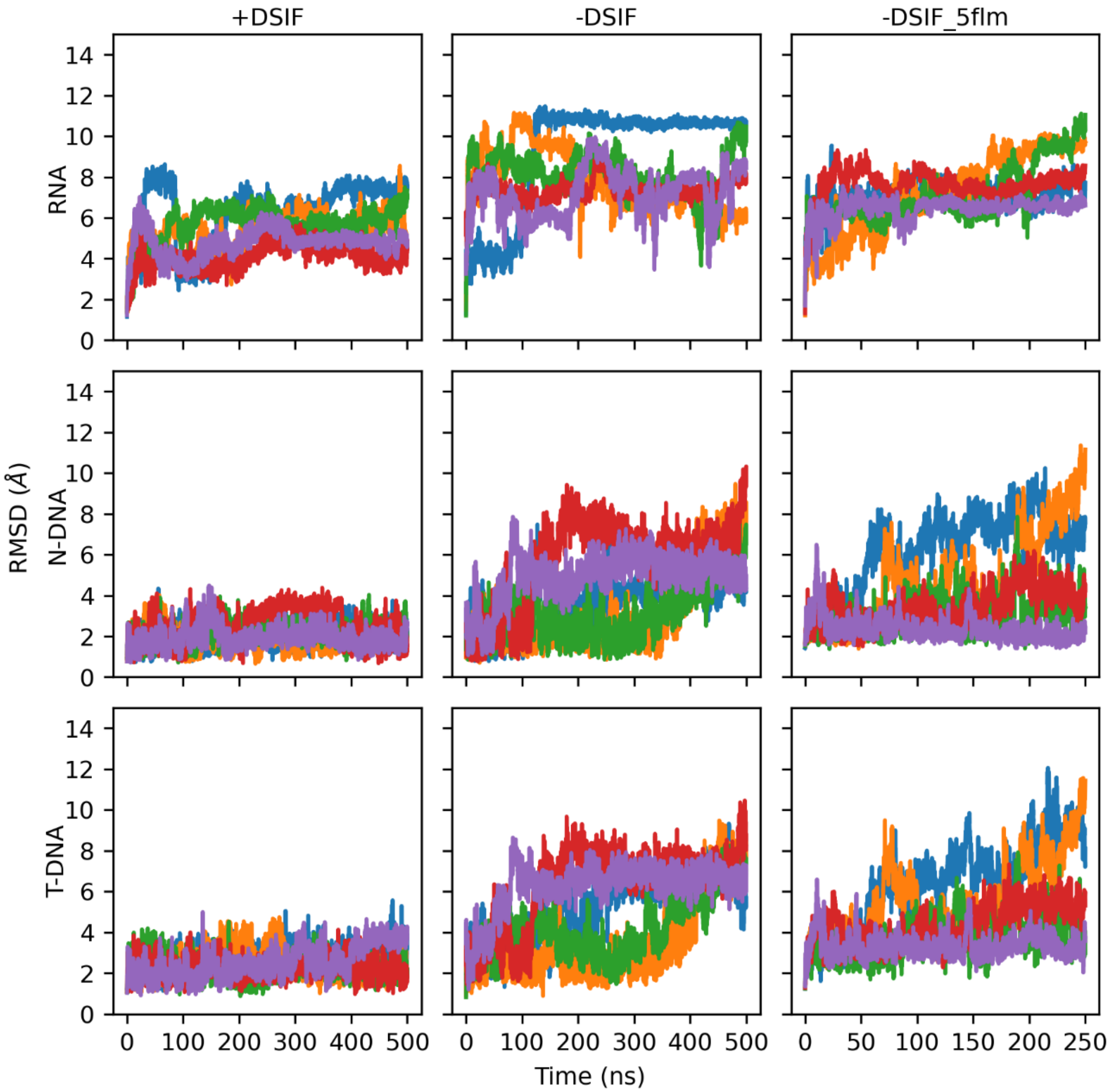


**Figure S2.** RMSD of the nascent RNA, upstream non-template (N-DNA) and template (T-DNA) DNA for the systems +DSIF, -DSIF and -DSIF_5flm. Lines with different colors represent RMSD from each replicate simulation.

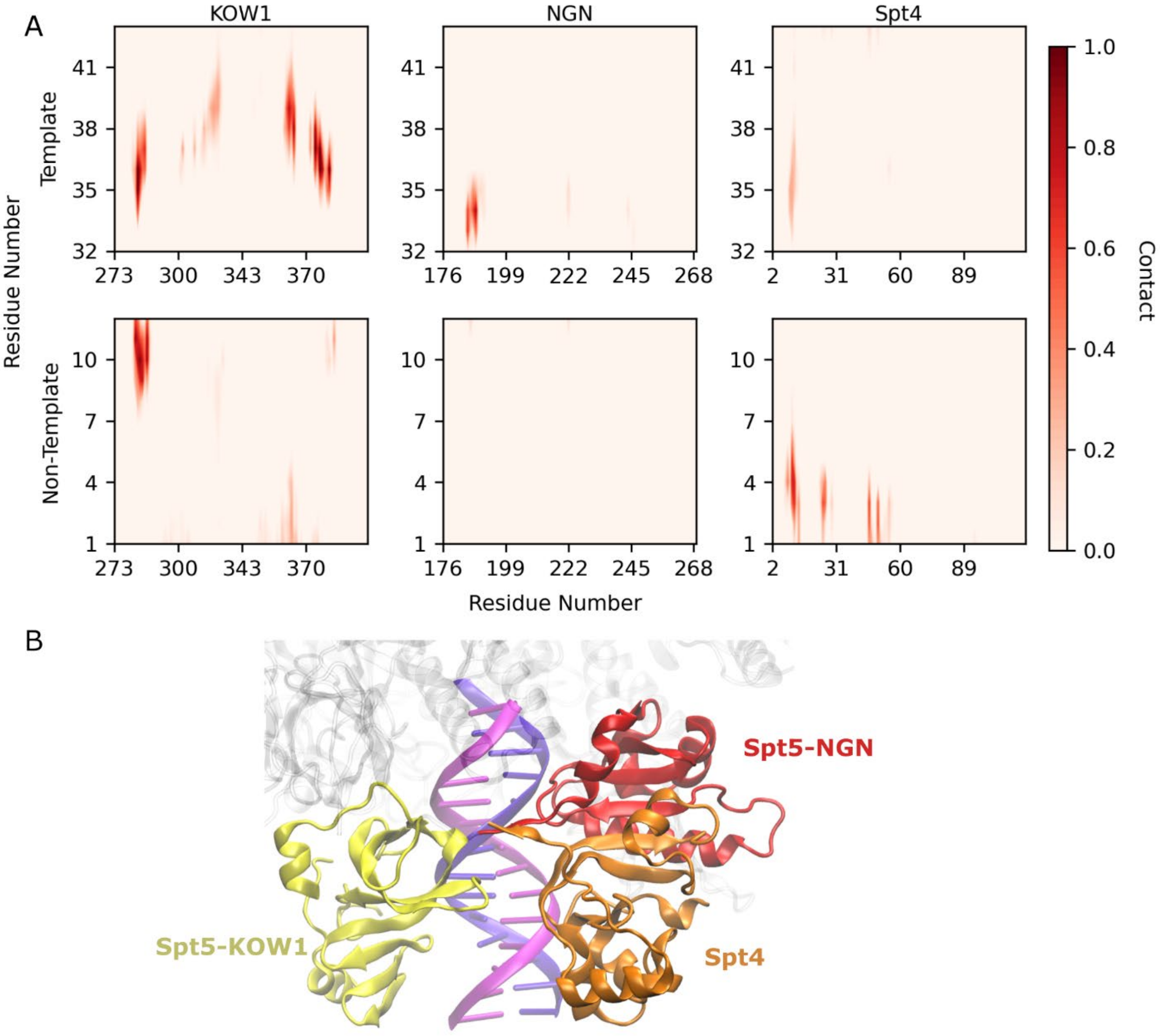


**Figure S3.** Contact maps for template and non-template upstream DNA with Spt5-KOW1, NGN domains, and Spt4 (A). Structure of upstream DNA surrounded by DSIF (Spt5-KOW1, NGN domains, and Spt4), shown in the initial structure (B).

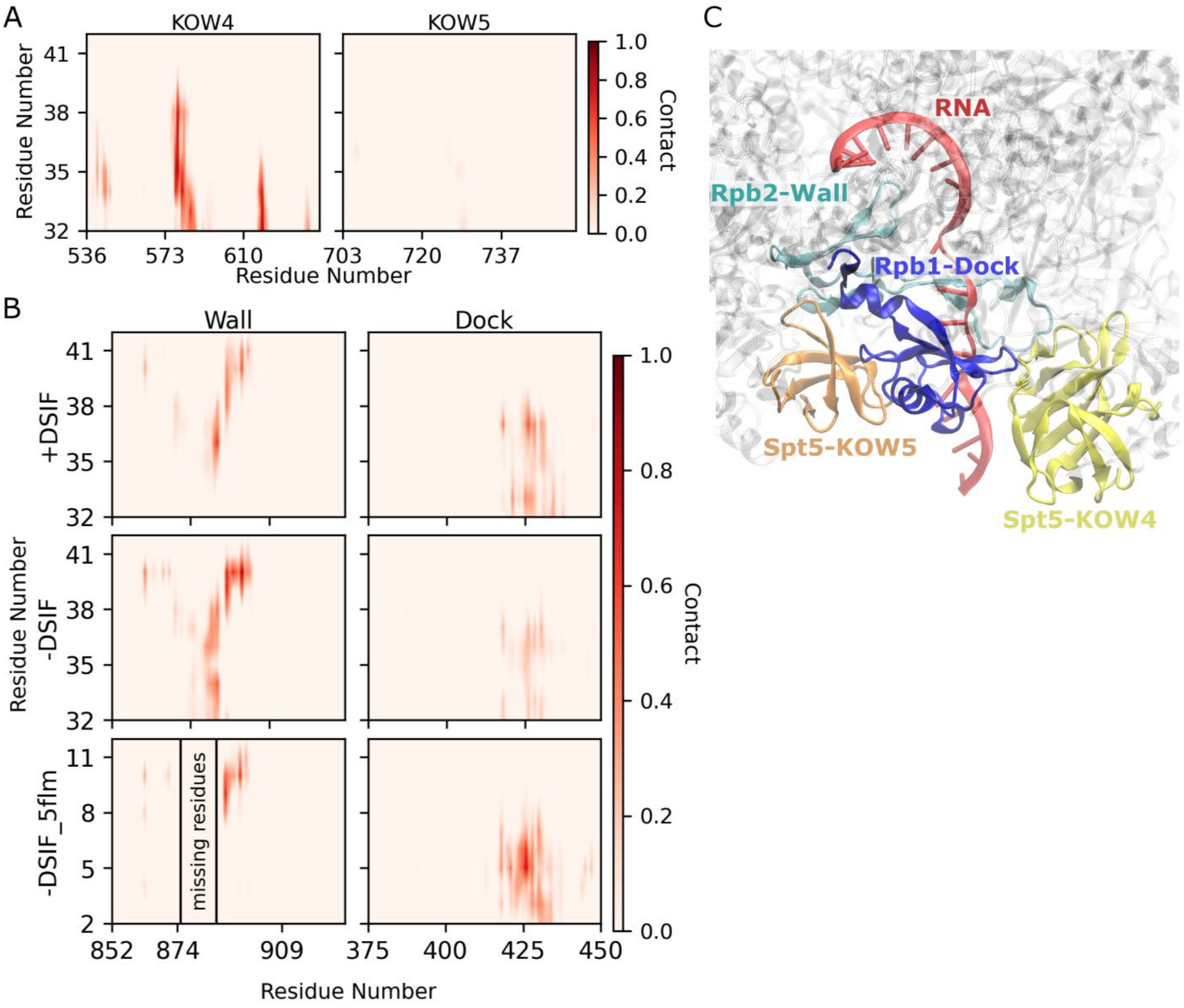


**Figure S4.** Contact maps for RNA with Spt5-KOW4 and KOW5 domains (A), and Rpb2-wall and Rpb1-dock domains (B). Structure of RNA surrounded by DSIF (Spt5-KOW4 and KOW5 domains), Rpb1 (dock domain) and Rpb2 (wall domain) shown in the initial structure (C).

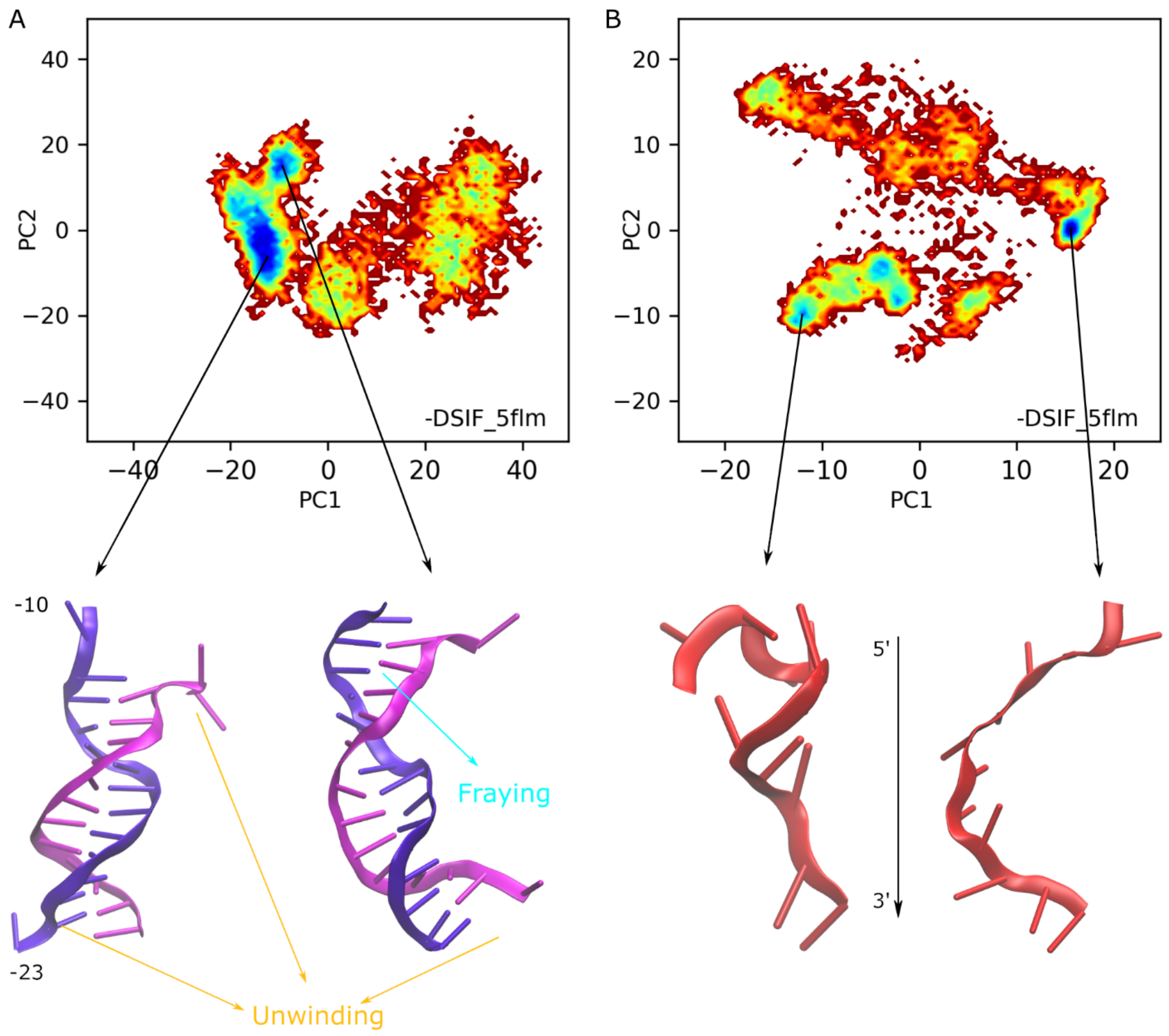


**Figure S5.** Free energy profiles for the conformations of the upstream DNA (A) and nascent RNA (B). The most populated conformations for the two lowest energy regions were provided at the bottom of each energy profiles.

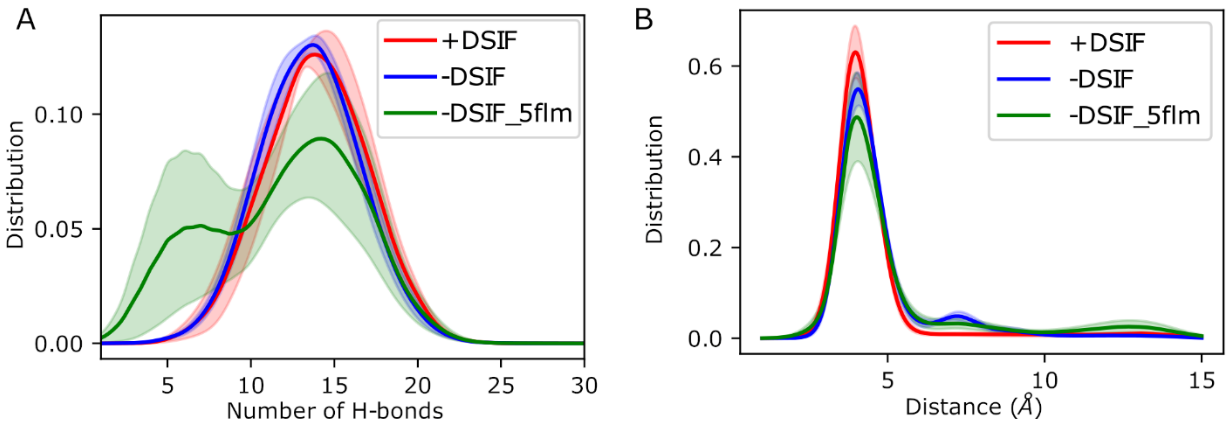


**Figure S6.** Distributions for the number of Watson-Crick hydrogen bonds (A) and base stacking distances (B) for the upstream DNA.

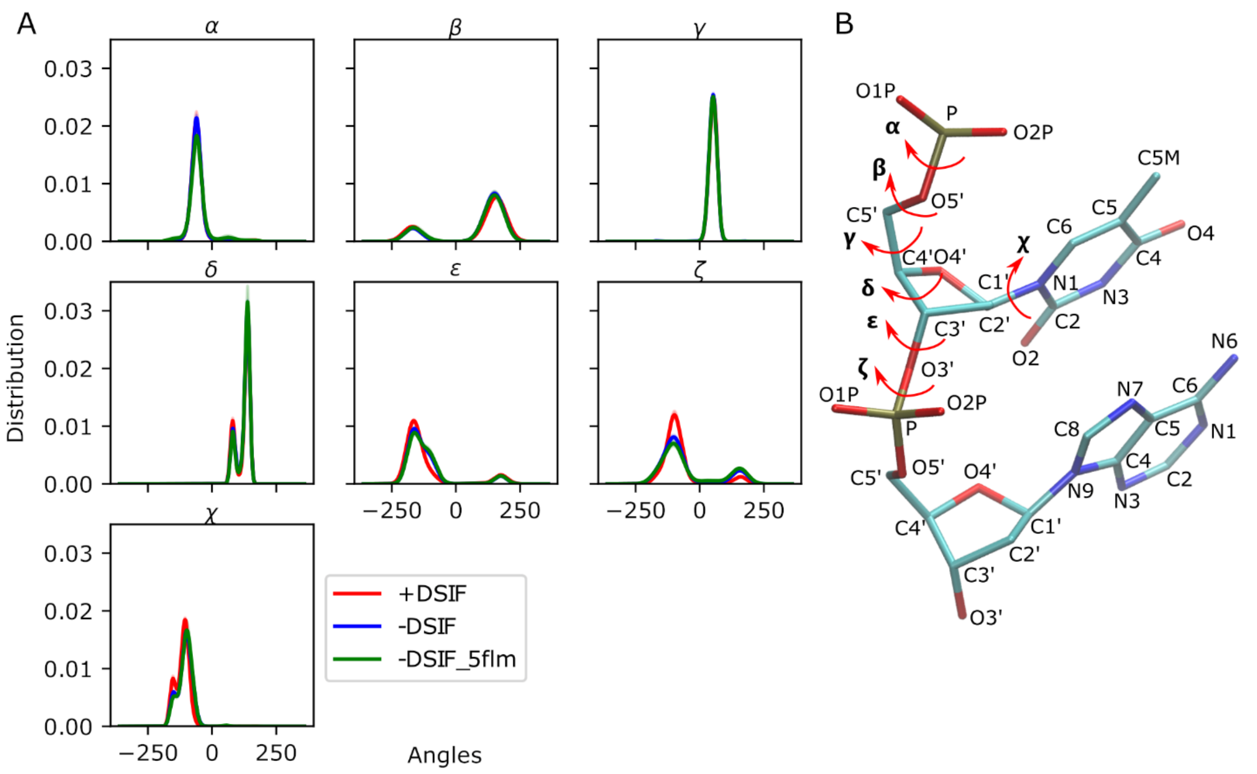


**Figure S7.** Distributions of dihedral angles for the upstream DNA (A), structural depictions of dihedral angles of nucleic acids (B).

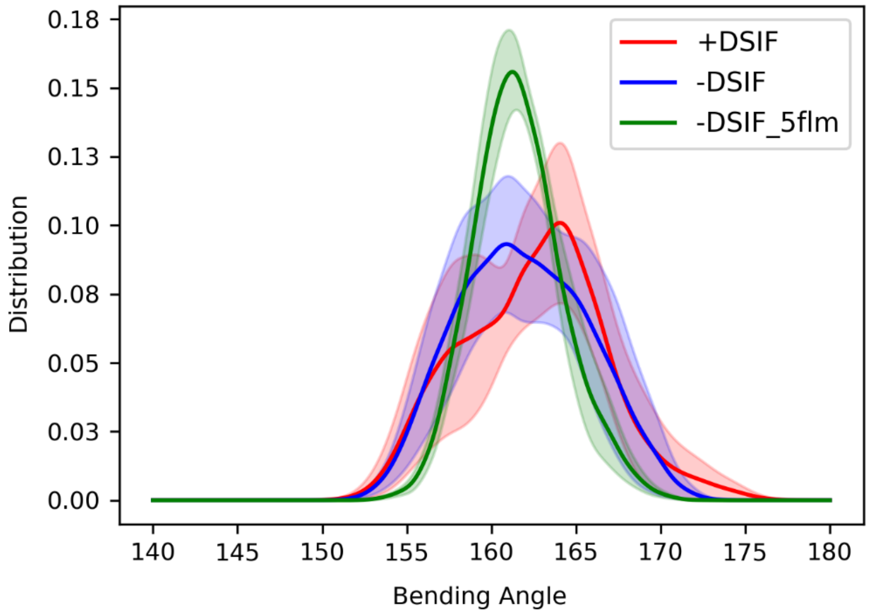


**Figure S8.** Distributions of the bending angles of the BH domain. Bending angles were calculated as the angle between the principal axes of two helices made by the residues 835-852 and 853-870 (see Methods for details).

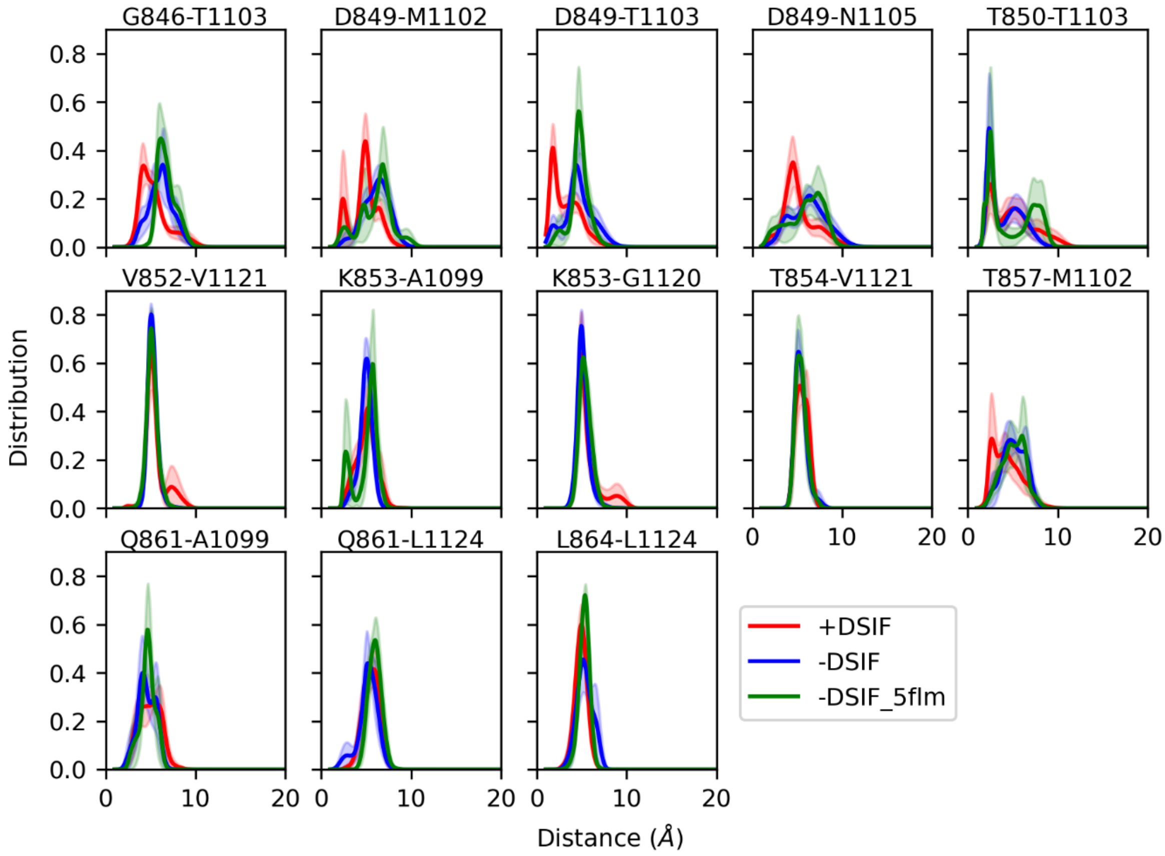


**Figure S9.** Distribution of distances for selected residue pairs of BH and TL. The pairs that have normalized contact more than 0.7 in either +DSIF or -DSIF systems, with a difference of 0.1 in normalized contacts between the two systems, were selected.

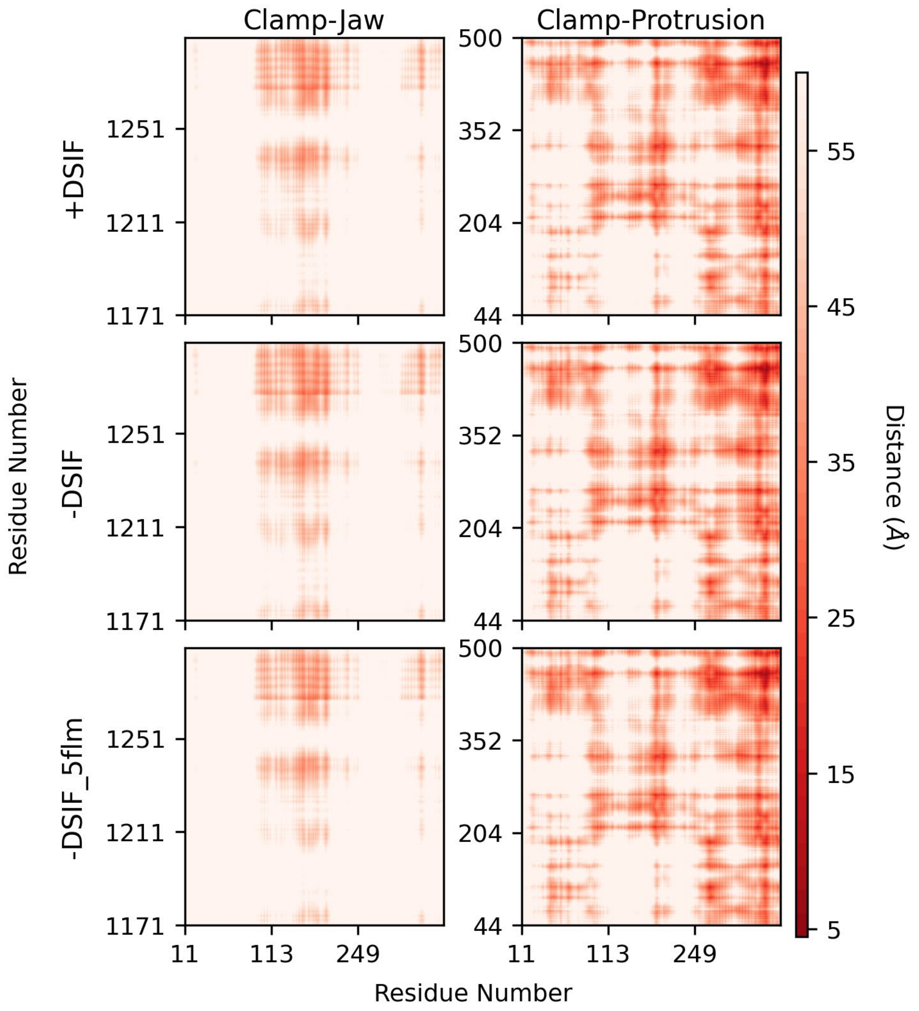


**Figure S10.** Distance maps for Rpb1-clamp and Rpb1-jaw, and Rpb1-clamp and Rpb2-protrusion.

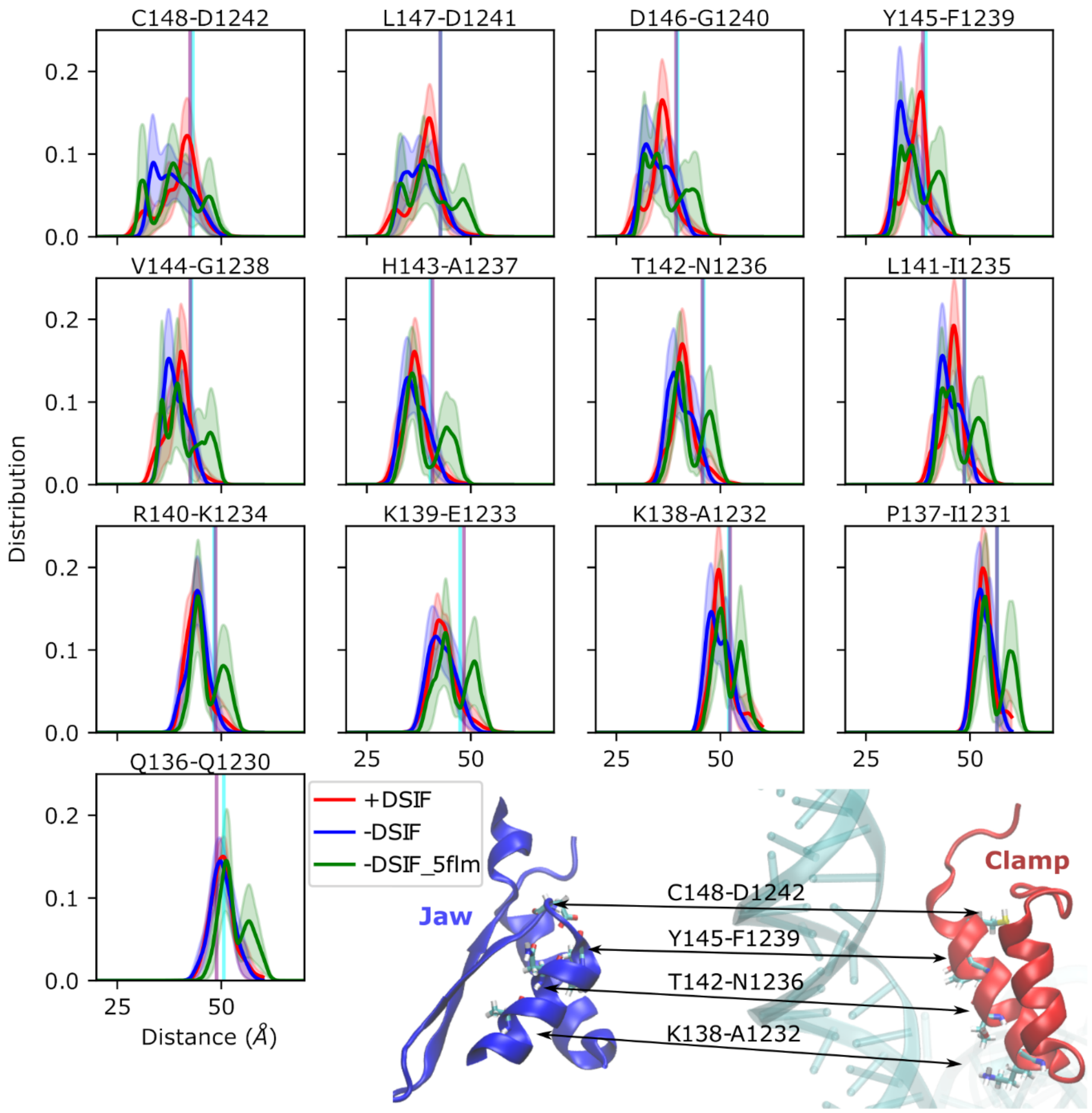


**Figure S11.** Distribution of distances for selected residue pairs of Rpb1-clamp and Rpb1-jaw. The pairs being parallel along each helix in two domains were selected, and examples of the four pairs are shown in the structure at the bottom of the figure. The vertical lines in cyan and purple show the values for the initial structures for PDB IDs 5FLM and 5OIK, respectively.

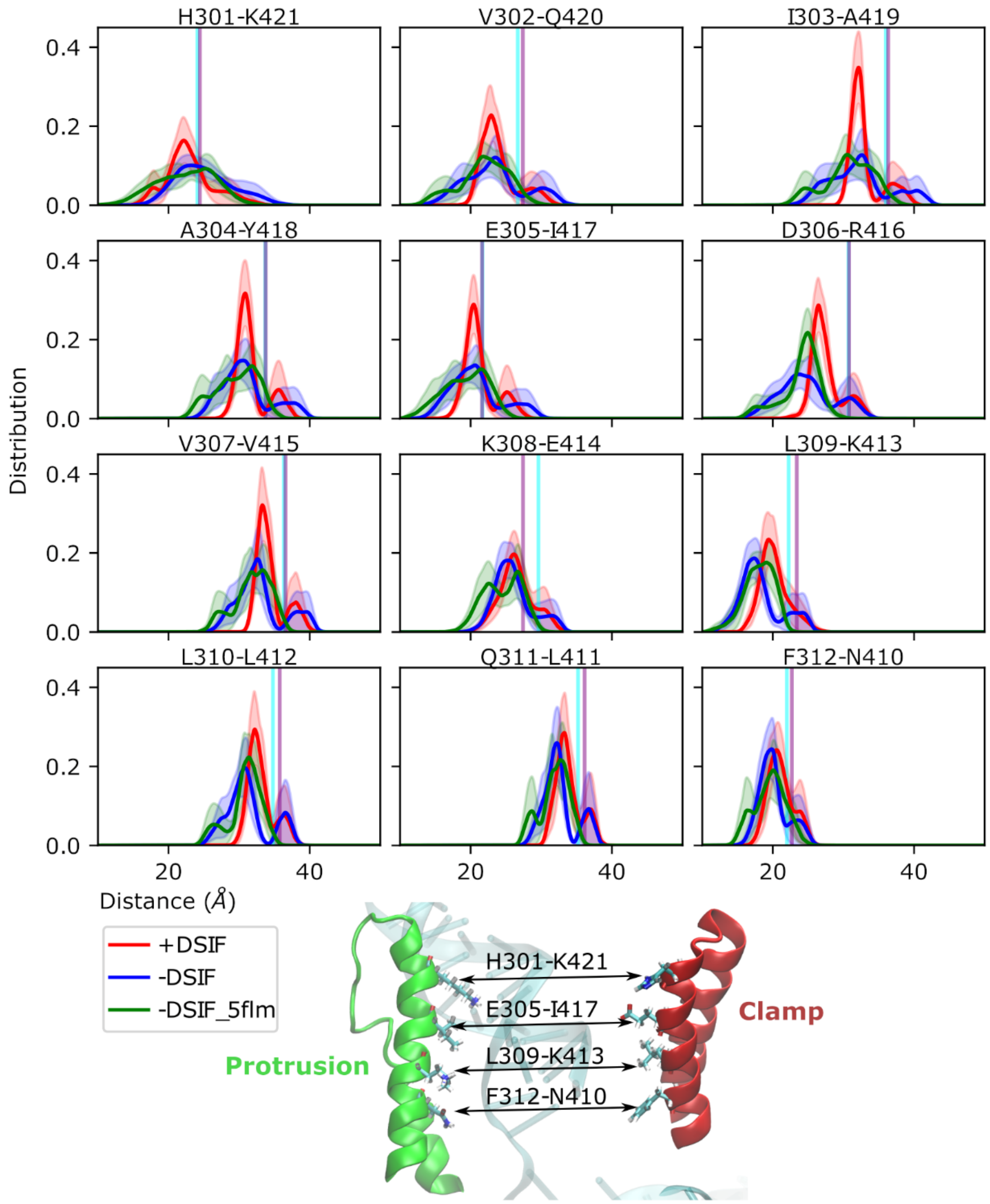


**Figure S12.** Distribution of distances for selected residue pairs of Rpb1-clamp and Rpb2-protrusion. The pairs being parallel along each helix in two domains were selected, and examples of the four pairs are shown in the structure at the bottom of the figure. The vertical lines in cyan and purple show the values for the initial structures for PDB IDs 5FLM and 5OIK, respectively.

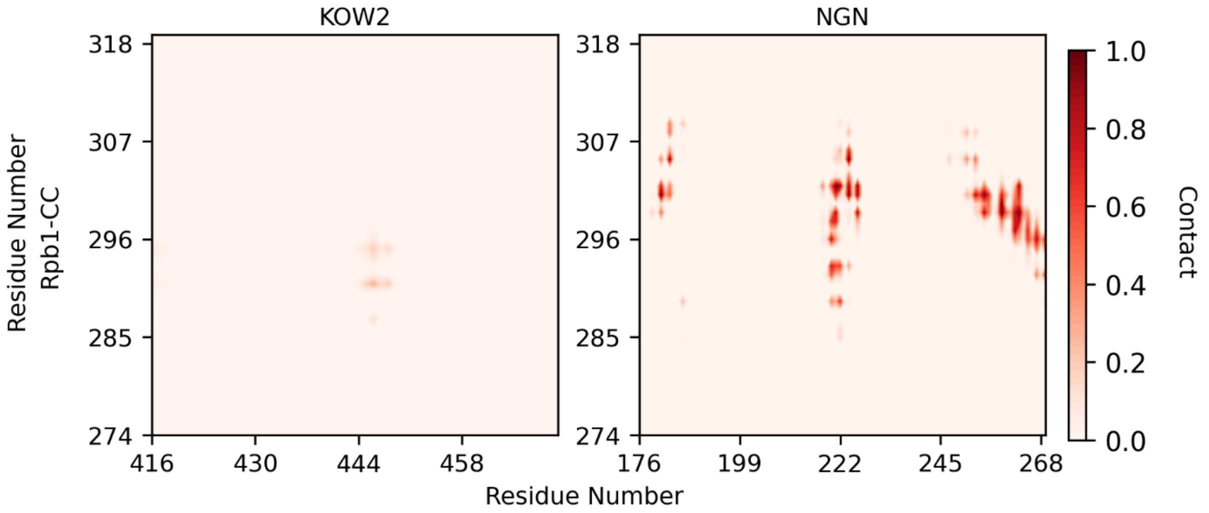


**Figure S13.** Normalized number of contacts for Rpb1-coiled coil (CC) domain with Spt5-KOW2 and NGN domains.

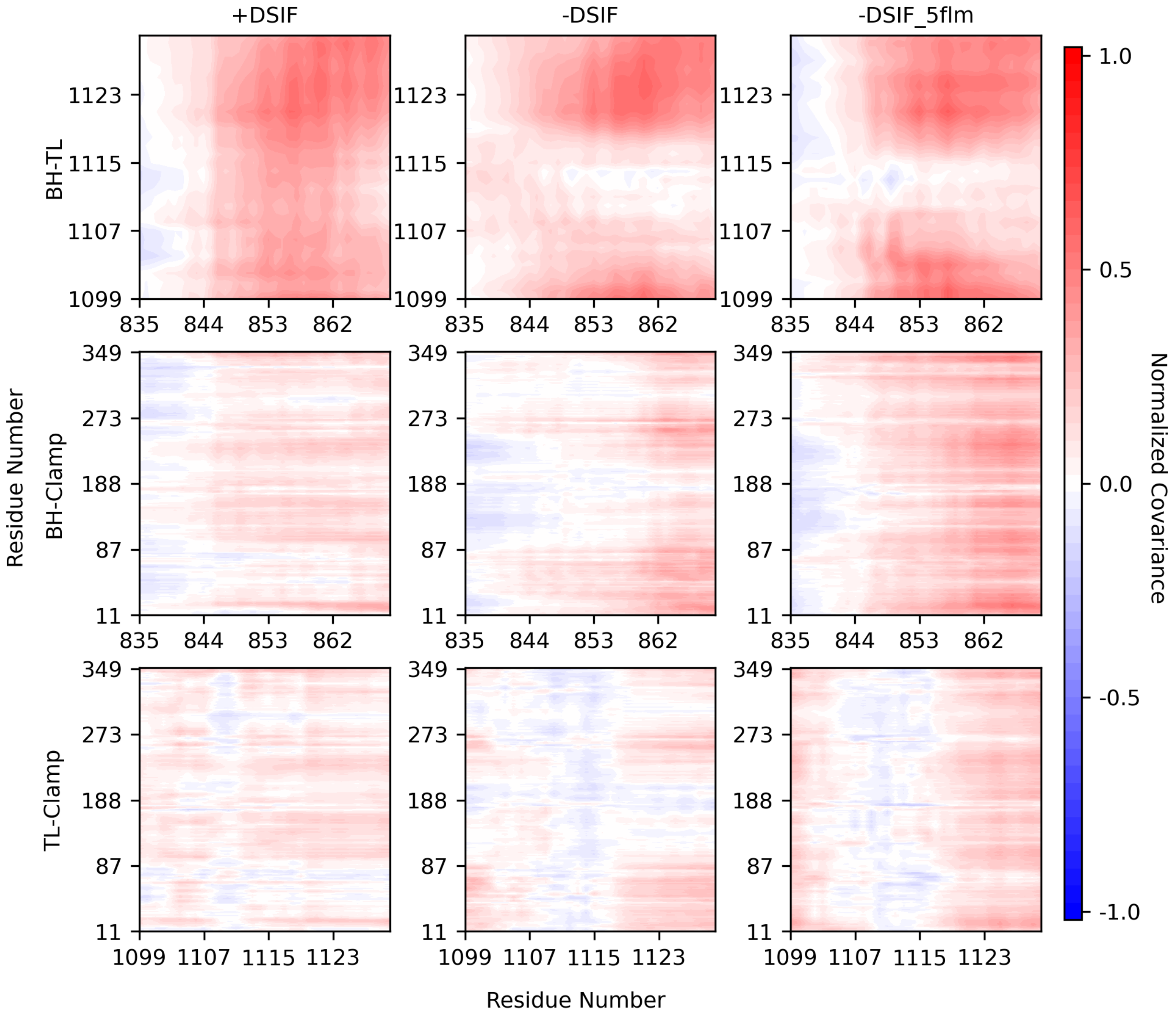


**Figure S14.** Cross-correlation maps between BH and TL, BH and clamp, and TL and clamp.

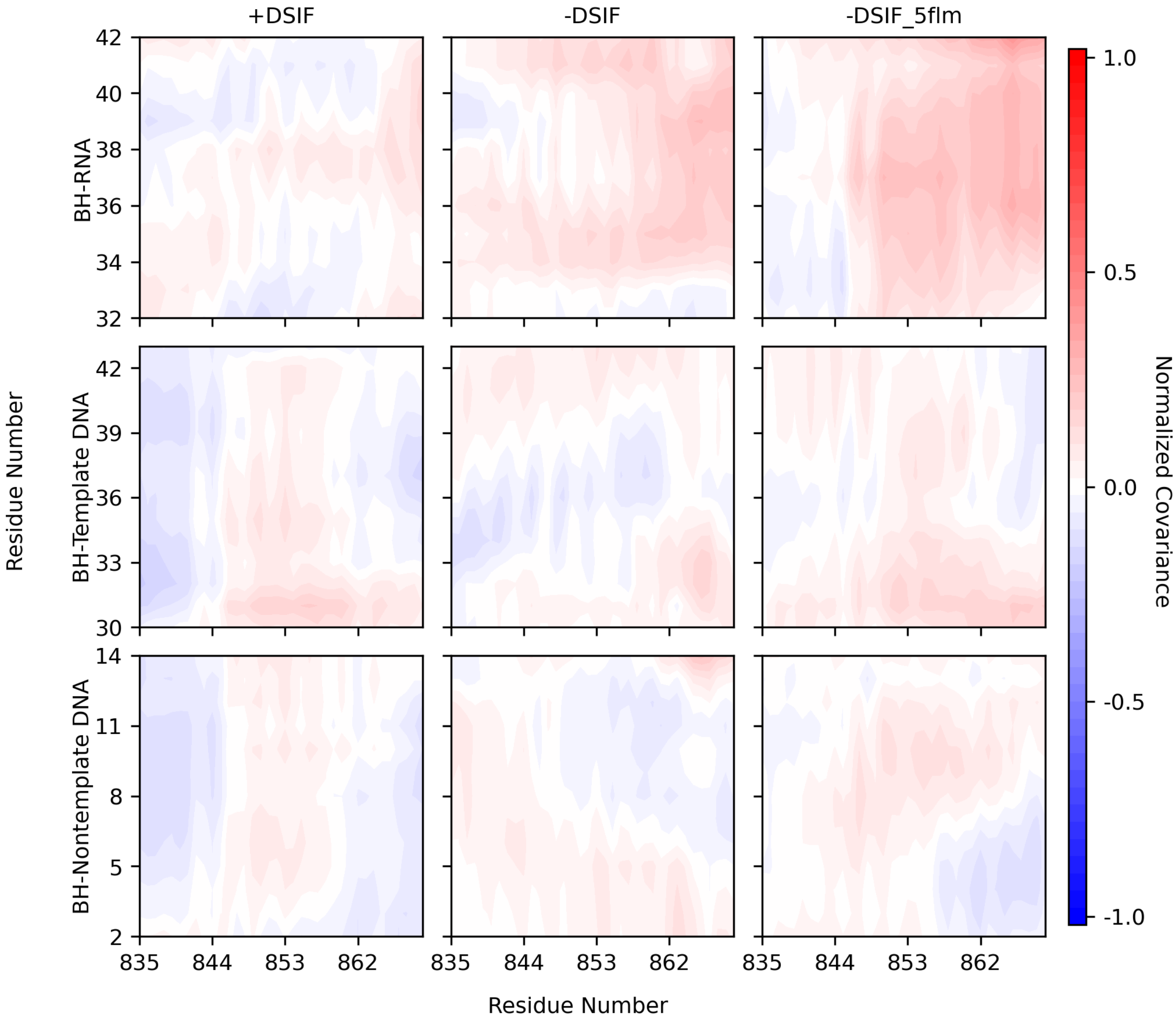


**Figure S15.** Cross-correlation maps between BH and RNA, BH and template, and non-template DNA.

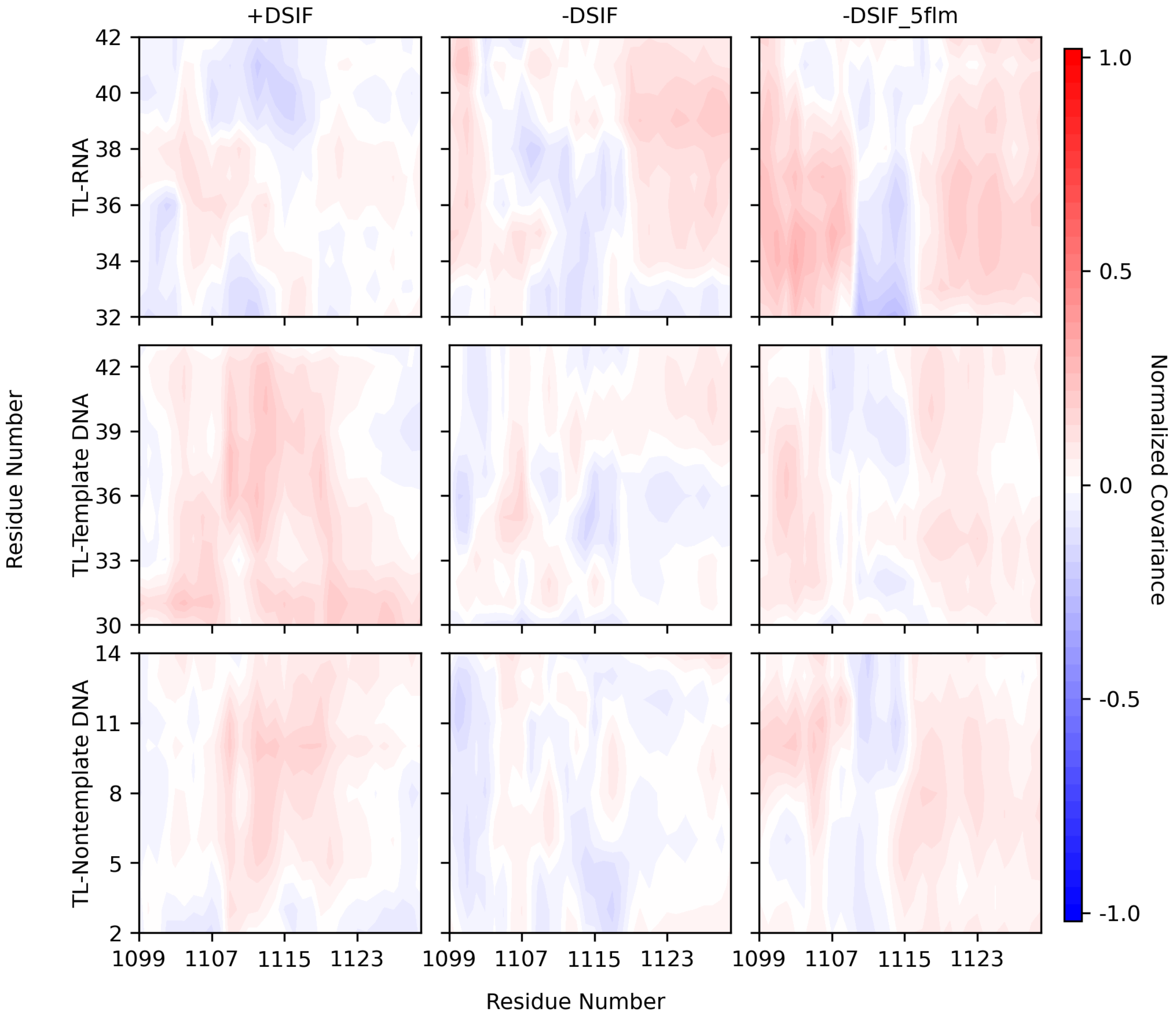


**Figure S16.** Cross-correlation maps between TL and RNA, TL and template, and non-template DNA.

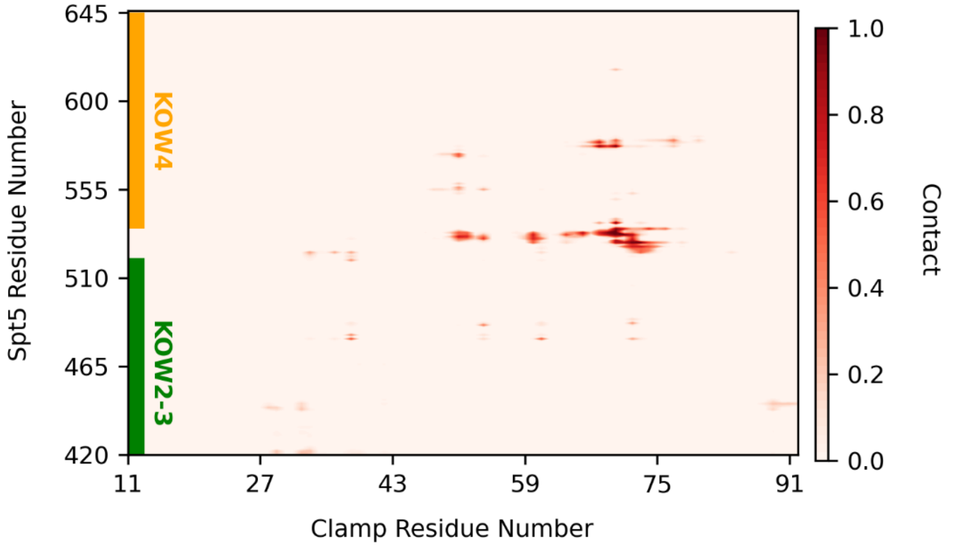


**Figure S17.** Normalized number of contacts for the residues 11-92 of the Rpb1-clamp domain with Spt5-KOW2-4.

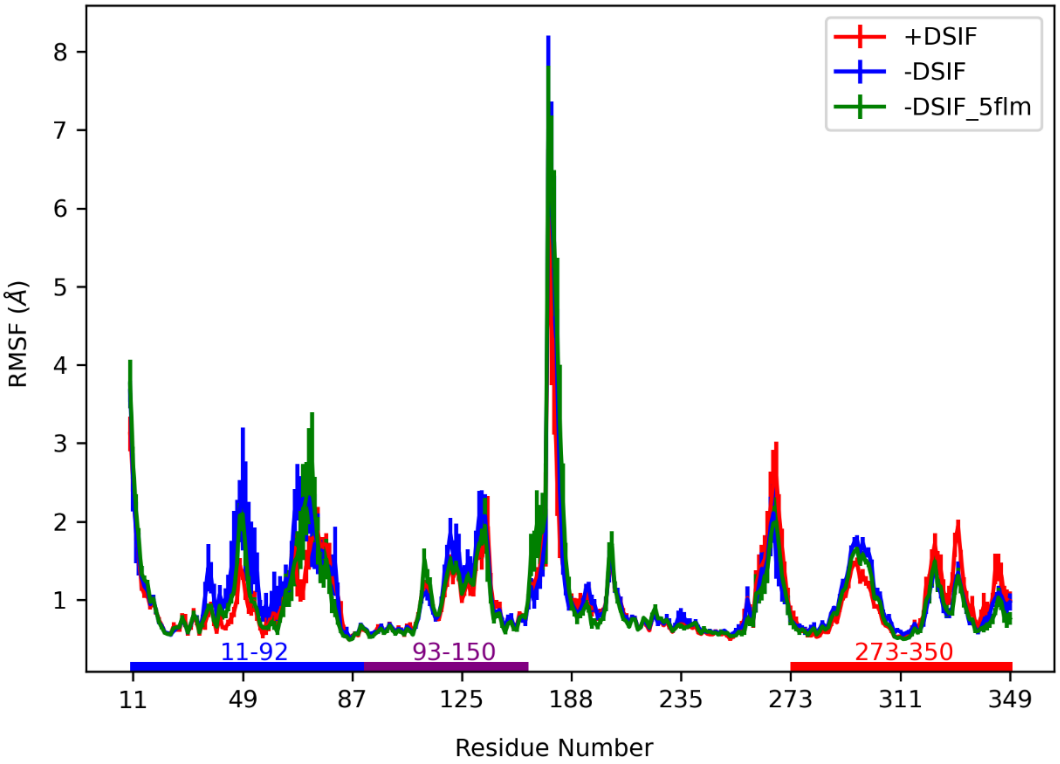


**Figure S18.** The root mean square fluctuations (RMSF) for Rpb1-clamp. The residues 11-92, 93-150, and 273-350 correspond to the same region highlighted with the same colors in Fig. 4B of the main manuscript.

**Table S1.** RMSD values between the structures with PDB IDs 5OIK and 5FLM.

| | RMSD fit to the selection | RMSD fit to Rpb1 | RMSD fit to Rpb1+Rpb2 |
|---|---|---|---|
| Rpb1 | 0.58 | 0.58 | 0.59 |
| TL original | 0.34 | 0.39 | 0.41 |
| TL modeled | 2.02 | 2.19 | 2.18 |
| BH | 0.26 | 0.29 | 0.28 |
| Rpb1-clamp | 0.72 | 0.73 | 0.75 |
| Rpb1-jaw | 0.19 | 0.21 | 0.21 |
| Rpb2-protrusion | 0.25 | 0.38 | 0.27 |

RMSD values were calculated for selected domains for three different types of fitting and provided in Å. For TL residues RMSD were calculated for the residues between 1099 and 1130 (TL modeled), which were partly modeled for the two systems and for the residues of two regions in the original structures (TL original), which the residues are 1099-1102 and 1115-1130.
